# A winding road to coexistence: Interdependence of niche and fitness differences in *E. coli* with targeted resource uptake gene deletions

**DOI:** 10.64898/2026.06.26.734880

**Authors:** Brendon McGuinness, Frederic Guichard, Stephanie C. Weber

## Abstract

Resource competition theory typically assumes static traits and continuous supply of resources. Yet microbial communities often experience feast–famine cycles and rapid trait change. To investigate coexistence under these nonequilibrium conditions, we integrate modern coexistence theory (MCT) with a genome-scale metabolic model that explicitly links resource use (traits) to metabolic fluxes and growth. MCT partitions competitive interactions into niche and fitness differences, to predict when trait-driven departures from neutrality result in coexistence or exclusion. Using dynamic flux balance analysis, we define a function that maps trait–resource matching to niche and fitness differences between species in a two-species two-resource system. This mapping shows that niche and fitness differences are not independently tunable under resource competition: changes in transporter-mediated resource uptake and changes in resource concentration ratios generate constrained trajectories through coexistence space. Specifically, we show that the minimum niche difference required for coexistence increases linearly with the absolute difference in maximal growth rates on limiting resources, showing how limiting similarity between species can emerge from intracellular metabolic constraints. Furthermore, we find that in batch culture simulations, initial conditions (inoculum size, total resource concentration) determine the timescale of the transient growth phase, with niche differences saturating and fitness differences increasing as the timescale grows, thereby governing competition outcomes. Finally, we test these predictions experimentally using *E. coli* strains with targeted resource transporter knockouts under both equal and skewed resource concentrations. Our results confirm that transporter-mediated trait changes and resource concentration ratio modulation can be harnessed to engineer coexistence. Together, our work demonstrates that trait-resource matching imposes structured constraints on the joint evolution of niche and fitness differences, thereby shaping biodiversity maintenance in microbial communities under nonequilibrium conditions.

---

A fundamental goal in ecology is to understand how diverse communities persist despite competition for similar resources. Classical coexistence theory posits that coexistence is maintained through stable niche partitioning, whereby species limit their own growth more strongly than that of their competitors, allowing multiple species to persist stably at equilibrium [1, 2, 3]. Modern coexistence theory (MCT) predicts coexistence by its decomposition into two independent components: niche differences (ND) that stabilize coexistence through negative frequency dependence and fitness differences (FD) whose reduction promotes coexistence through fitness equalization [4, 5, 6]. However, recent work challenges the independence of stabilization and equalization, instead suggesting their covariance under the action of specific mechanisms such as resource competition [7]. Mechanistic consumer-resourcemodels reveal that even small changes in traits (corresponding to differences in resource use) often change both niche and fitness differences simultaneously, constraining the trajectories of competing species on the plane defined by axes of niche and fitness differences.

In the limiting case of neutrality, there are no niche or fitness differences between ecologically equivalent species [8, 7]. Small trait changes could correspond to speciation through niche differentiation associated with the loss of ecological equivalence [9], or could be due to intraspecific phenotypic variation [10, 11, 12]. The specific trajectories in the ND-FD plane on which competing species leave neutrality can lead to stable coexistence if variations in niche and fitness differences are independent. However, they can also lead to exclusion if trait changes lead to co-varying changes in ND and FD. This interdependence between niche and fitness differences can thus predict the minimum niche differentiation, or limiting similarity, required to overcome associated changes in fitness inequality and lead to stable coexistence [1, 13, 7].

In microbial systems, multiple mechanisms could mediate the interdependence between niche and fitness differences. Resource-use plasticity via gene regulation, de novo mutation, and horizontal gene transfer can alter metabolism within a few generations [14, 15, 16, 17] and influence both niche and fitness differences[7, 17]. Departures from neutrality are thus unlikely to move along a single ‘niche’ or ‘fitness’ axis and trajectories through coexistence space are expected to be constrained. Crucially, these rapid trait changes unfold in far-from-equilibrium environments, where pulsed resource supply and recurrent bottlenecks may shape coexistence outcomes [18]. Microbial communities often experience nonequilibrium feast-famine cycles such as in the gut microbiome [19], wastewater treatment [20], and particle associated marine snow [21]. These conditions can be imitated in serial-dilution batch cultures that contrast sharply with the steady resource supply of chemostats [22, 23, 24]. Explaining and predicting coexistence under such feast-famine regimes requires theoretical frameworks that link measurable traits and resource environments to the coupled evolution of niche and fitness differences under nonequilibrium dynamics.

Recent advances in systems biology have produced genome-scale metabolic models that integrate vast genomic and biochemical knowledge into predictive computational frameworks [25, 26]. Dynamic flux balance analysis (dFBA) frameworks, such as COMETS, use these metabolic network reconstructions to simulate population growth, metabolite exchange, and resource dynamics by optimizing intracellular fluxes subject to stoichiometric constraints [27, 28]. Despite their common use in microbial ecology and metabolic engineering, dFBA has not yet been integrated into MCT. Bridging these domains would ground niche and fitness differences in measurable metabolic traits and species–environment feedbacks. Such an integration would move coexistence theory toward mechanistic explanations that are testable in highly controlled microcosm experiments, and thereby narrow the gap between rapid empirical advances and our theoretical understanding of microbial community dynamics.

Building on theoretical work that shows that mechanistic changes in resource uptake can jointly shift niche and fitness differences [7], we reason that dFBA can be used to map constrained departures from neutrality in microbial competition assays: how altering a specific uptake trait and resource concentration ratios together channel communities through coexistence space. Linking such mechanistic trajectories to MCT predictions would bridge metabolic and coexistence perspectives, revealing general principles by which metabolic constraints and trait evolution govern the emergence and maintenance of biodiversity. Here, we apply this approach to pairs of *E. coli* strains competing for pairs of carbon sources under batch-culture conditions (i.e. feast-famine). Using COMETS dFBA, we define a departure-from-neutrality function that couples a single transport-flux trait (maximum uptake rate on the alternative carbon) with relative resource concentrations and identities. Sampling 36 carbon source-pair combinations, we trace vector trajectories through coexistence space and show that resources with large maximal growth rate differences impose a limiting-similarity threshold, where greater niche differentiation is required to balance fitness asymmetries. We then engineer transporter-knockout *E. coli* strains to modulate uptake asymmetries and perform reciprocal invasion assays under equal and skewed glucose–succinate supplies to test model predictions. Thus, by coupling metabolic modeling on a genome scale with controlled invasion experiments, we show that metabolic and environmental constraints shape the joint evolution of niche and fitness differences, providing a tractable framework to predict and engineer coexistence in microbial communities.

## Results

### A metabolic mapping of resource competition onto niche–fitness space

To determine how competition for substitutable carbon sources structures coexistence outcomes, we instantiated a metabolic framework using COMETS dFBA [28]. Two otherwise isogenic strains of *E. coli* supplied with two carbon sources differ only in a trait *T*, which represents the uptake flux of the alternative (less preferred) carbon source: strain 1 carries a reduced uptake bound (*T* ≤ *T*_*max*_) on carbon source *β* while strain 2 carries the same reduction on carbon source *γ*, producing a symmetric flux matrix that controls niche overlap (Figure 1A).

**Figure 1.**
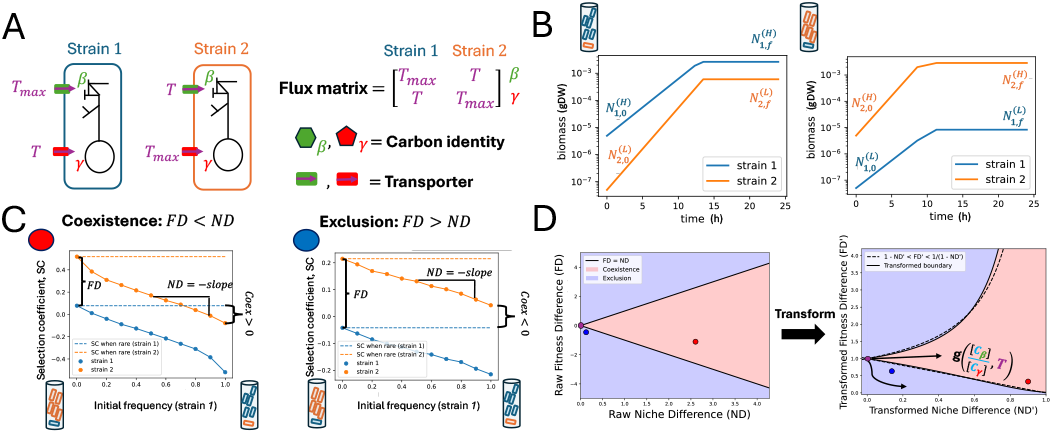
Metabolic mapping of resource competition onto niche–fitness space using dynamic flux balance analysis (dFBA). **(A)** Schematic of parameters that affect coexistence. Strains 1 and 2 represent nearly identical strains of *E. coli* in a genome-scale metabolic model. They differ only in *T*, the maximum uptake flux of carbon resource *β, γ* from outside to inside of the cell. Flux matrix of strains 1 and 2 in an environment with carbon sources *β* (denoted by green hexagon) and *γ* (denoted by red pentagon). The green and red boxes correspond to the transporters for these carbon sources, respectively. **(B)** Example of reciprocal invasion assay where strain 1 is in blue and strain 2 is in orange. 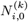 represents the initial biomass (grams dry weight GDW) at (*t* = 0) of strain *i* when it starts *k* ∈ (*L, H*) with where *L* is low initial biomass (invader) and *H* is high initial biomass (resident). Biomass time-series output of dFBA where N represents biomass of strain 1 (blue) or 2 (orange) at either initial or final time points and high (resident) or low (invader) initial frequency. **(C)** Two examples visualizing our method of measuring niche difference (*ND*), fitness difference (*FD*), and coexistence strength (*Coex*) from dFBA. The scenario on the left corresponds to coexistence (*T* = 1, *T*_*max*_ = 10, *β* =glucose, *γ* =citrate) and the scenario on the right to exclusion (*T* = 6, *T*_*max*_ = 10, *β* =glucose, *γ* =citrate). **(D)** On the left hand side is a raw coexistence phase diagram with the coexistence region, where |*FD*| *<* |*ND*|, shaded red and the exclusion region, where |*FD*| *>* |*ND*|, shaded blue. The boundary conditions, where *FD* = *ND*, corresponds to when Coex= 0 (Eq. 9). On the right hand side is the transformed coexistence phase space, in which the transformed niche and fitness differences from our method are defined as 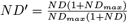 and *FD*^*′*^ = *e*^*FD*^, respectively (Eq. 10-11). The dashed lines represent the boundary conditions from Eq. 3, where 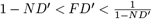_*′*_ to the scenarios in (C), where the red dot corresponds to the coexistence case (left side) and the blue dot corresponds to the exclusion case (right side). The purple dot corresponds to neutrality, i.e. *T* = *T*_*max*_. The black arrows leaving neutrality represent potential constrained trajectories which are represented by the function 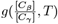 and lead to different competitive outcomes.

We quantified competitive outcomes through reciprocal invasion assays, in which each strain was initialized at high (resident) and low (invader) biomass against its competitor (Figure 1B). From the resulting biomass time series, we computed selection coefficients for each strain under various initial frequencies (Methods). The niche difference (*ND*) was estimated as the strength of negative frequency dependence, the fitness difference (*FD*) as the difference in selection coefficients when rare, and coexistence strength (*Coex*) as the minimum selection coefficient when rare (Figure 1C; Eqs. 7–9 in Methods). Mutual invasibility and thus coexistence is predicted when coexistence strength is positive; exclusion occurs when one strain fails to increase from rarity. For example, a case with *T* = 1 in glucose and citrate yielded *FD < ND* and thus a positive coexistence strength (Figure 1C, Left), while *T* = 6 on the same resources produced *FD > ND* and thus exclusion (Figure 1C, Right). In both of these examples, *T*_*max*_ = 10.

To compare these simulation-derived metrics directly with analytical coexistence criteria from MCT, we transformed raw niche and fitness differences onto scales consistent with the Lotka–Volterra model (Figure 1D; Methods). The transformed metrics (*ND*^*′*^, *FD*^*′*^) aligned closely with theoretical coexistence boundaries (Eq. 3), confirming that invasion-derived quantities from dFBA simulations preserve the predictive structure of MCT in its Lotka-Volterra-derived form (Figure 1D). Under this mapping, we defined a hypothetical constraint function, 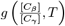, in which shared resource use does not generate arbitrary combinations of niche and fitness differences, but instead channels departures from neutrality along structured trajectories toward coexistence or exclusion (Figure 1D). With our validated framework in hand, we next asked how the constraint function *g* structures trajectories through coexistence space across carbon source pairs that differ in identity, relative concentration, and quality.

### Maximum growth rate differences on carbon sources govern degree of limiting similarity

To investigate how departures from neutrality influence coexistence outcomes, we quantified the effects of varying resource-uptake on alternate carbon sources, *T*, and supplied carbon source concentration ratio,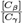, on trajectories through coexistence space. Starting from neutrality (*T* = *T*_*max*_ = 10), we traced vector trajectories for two different resource pairs: glucose and fructose (Figure 2A), as well as glucose and succinate (Figure 2B). Glucose and fructose are identical in quality (maximum growth rate and yield) in COMETS, leading to a weak effect of *T* on fitness differences when the relative concentrations of the supplied carbon sources were similar. With these resources, departures from neutrality led to stable coexistence without any minimum niche differentiation (limiting similarity) over a wide range of relative carbon source concentrations. However, as the relative resource concentration became very skewed, niche and fitness differences changed interdependently and exclusion occurred over much of the ND-FD trajectories. In the case of glucose and succinate (where glucose maintains a higher growth rate than succinate), we observed limiting similarity: regardless of the relative concentration of carbon sources (we tested over the range 1 : 10 − 10 : 1), initial niche differentiation through decreasing *T* imposed changes in fitness differences that led to exclusion until a threshold was reached below which *T* was small enough for coexistence to occur.

**Figure 2.**
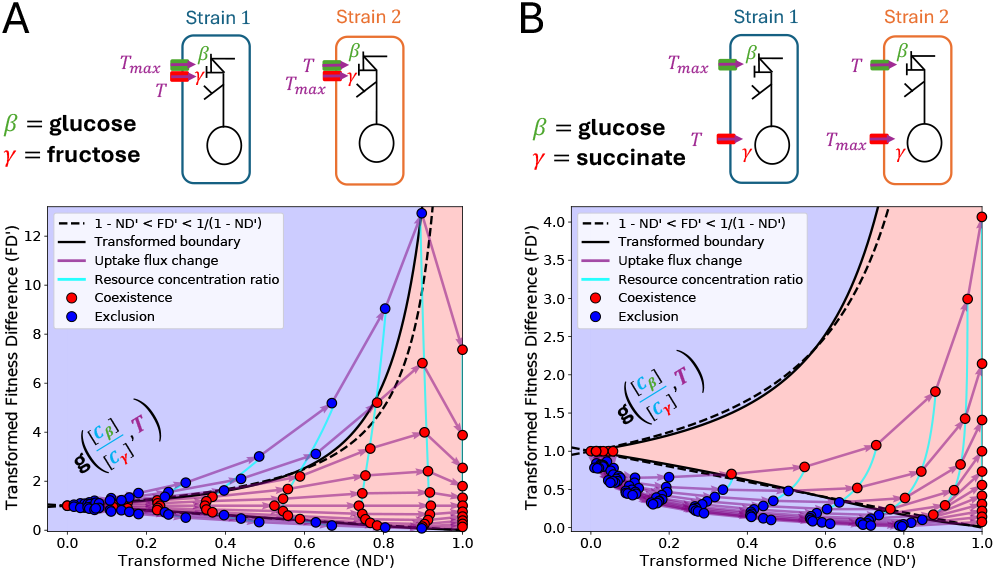
Departure from neutrality along constrained trajectories in coexistence phase space have different outcomes. **(A)** Diagram representing simulations of a co-culture departing from neutrality by changing the uptake flux, *T*, on alternate resources (*β* = glucose, *γ* = fructose) and the corresponding trajectories from neutrality by changing *T* (purple lines) or resource concentration ratio 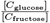 (cyan lines) from 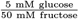 to 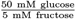. Axes correspond to transformed niche and fitness differences. **(B)** Diagram and trajectories from neutrality where *β* = glucose and *γ* = succinate by changing uptake flux on alternate resources (purple lines) or resource concentration ratio (cyan lines) as in (A).

Next, we evaluated 36 carbon source pairs at equal concentrations (27.5 mM) where *T* = 10, …, 0 and measured coexistence strength as a function of raw niche difference (Figure 3A). For every carbon source pair, we ran dFBA simulations of monocultures (with *T* = 10) on each individual carbon source and calculated the absolute difference in maximum growth rate. When carbon sources are equal or very similar in terms of nutrient quality (i.e. maximum growth rate difference is small), then coexistence was either always possible or constrained by a small limiting similarity. As maximum growth rate difference becomes larger due to differences in carbon source quality, the effect of limiting similarity became stronger and strains required larger niche differences to coexist (Figure 3B).

**Figure 3.**
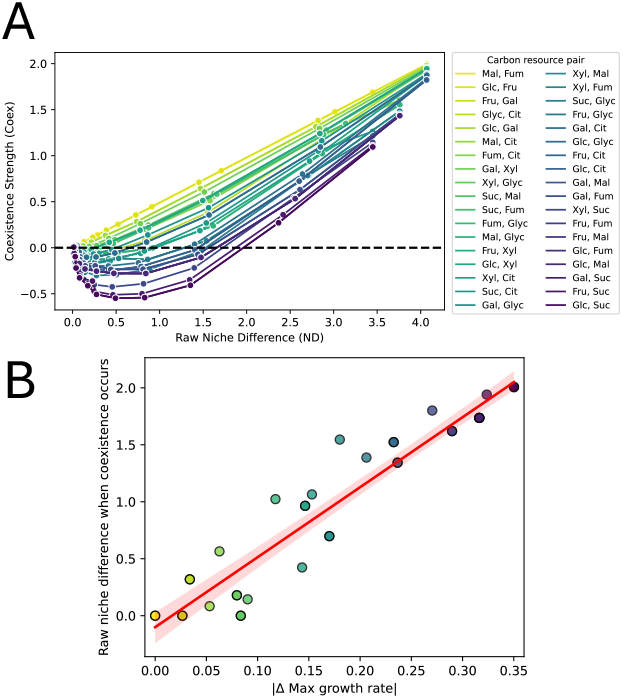
Degree of limiting similarity resulting from constrained vector fields along departure from neutrality depend on max growth rate differences on alternate resources. **(A)** Coexistence strength, Coex, as a function of raw niche differences for 36 pairs of carbon sources (*β, γ*) where concentrations are 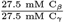. See Table S1 for details on carbon sources. Colormap corresponds to the absolute difference in max growth rate on single resource *β* vs. single resource *γ* of the wildtype strain where *T* = *T*_*max*_ = 10. Dashed line represents the coexistence boundary, where Coex= 0. **(B)** The minimum raw niche difference where coexistence occurs (Coex= 0) as a function of absolute difference in max growth on single resource *β* vs. single resource *γ*. The points are coloured corresponding to the legend in (A). Linear fit to data shown in red, with 95% confidence interval. Note that in (A) and (B) the raw niche differences were not transformed to preserve the linear relationship between (raw) niche difference at coexistence and absolute difference in max growth rate.

Since maximum growth rate difference predicts the degree of constraint on trajectories through the ND-FD phase space, we asked whether these growth-rate differences could be related to coarse biochemical features of individual carbon sources and their associated flux solutions. Across an extended panel of 23 single carbon sources, NADH production emerged as the strongest single correlate of maximum growth rate, while the number of active reactions provided additional explanatory power beyond NADH alone (Supplementary Text A; Figure S1). These results suggest that substrate-specific growth rates are not arbitrary outputs of the genome-scale metabolic model, but are associated with interpretable features of redox metabolism and network engagement. Thus, differences in substrate chemistry propagate through intracellular metabolism to generate the growth-rate asymmetries that help determine fitness differences and constrain trajectories through coexistence space.

Fitness differences are driven by maximal growth-rate asymmetries, whereas niche differences arise from frequency-dependent selection through resource partitioning. We therefore asked whether the duration of the exponential growth phase modulates their relative strength. We manipulated the timescale of this growth phase before the stationary phase (where the resources have run out) by changing the total initial concentration of resources as well as the initial biomass of the strains (Figure 4A,B). Increasing the total resources increased the timescale of this growth phase, and similarly, decreasing the initial biomass of the two species increased the timescale of the growth phase (Figure 4C). Increasing the timescale of the growth phase led to greater fitness differences but saturated niche differences (Figure 4D), thereby leading to a transition from coexistence to exclusion. These results show that transient growth duration regulates the balance between stabilizing and equalizing forces, strengthening limiting similarity as exponential growth is extended.

**Figure 4.**
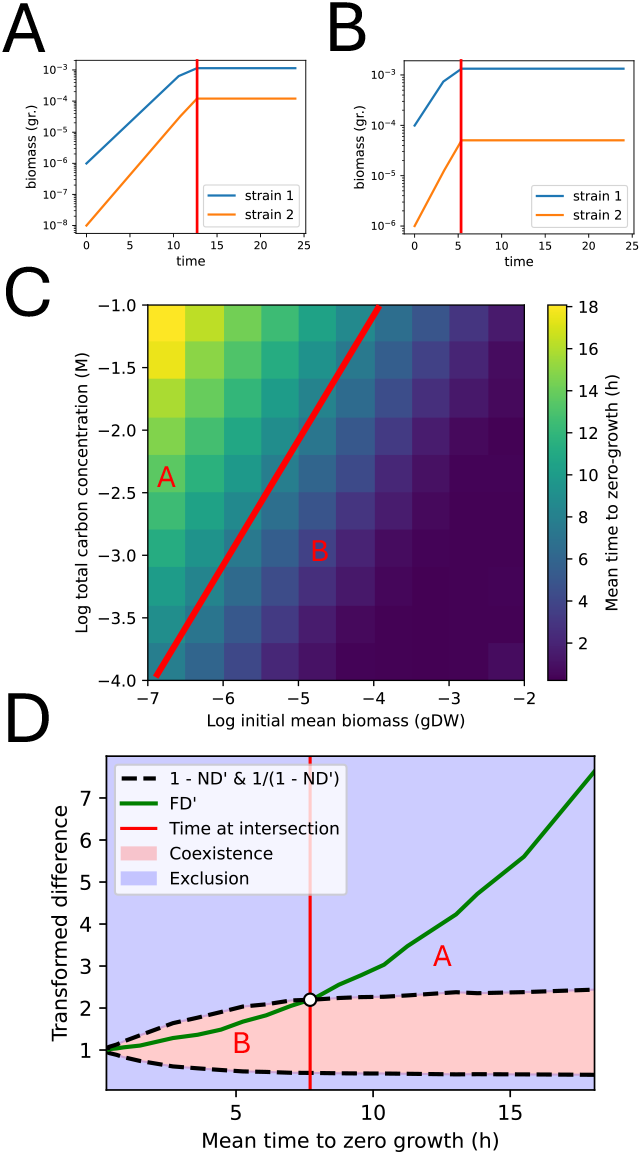
Competitive outcome depends on timescale of growth phase. **(A)** Time-series with low initial biomass and high total carbon source concentration that expands the growth phase. **(B)** Time-series with higher initial biomass and lower total carbon source that contracts the growth phase. **(C)** Heatmap showing the mean time to zero growth (duration of growth phase) as a function of both total carbon concentration and initial mean biomass. Red line corresponds to time of intersection point in (D) that separates coexistence and exclusion outcomes. The red letters correspond to the timescales, total carbon concentrations, and initial biomass values from (A) and (B), respectively. **(D)** Transformed niche (*ND*^*′*^) and fitness (*FD*^*′*^) differences as functions of the mean time to zero growth. The red line corresponds to the time when *ND*^*′*^ and *FD*^*′*^ intersect, and thus marks the transition from coexistence to exclusion. The red letters correspond to parameters from (A) and (B).

### Experimental validation using *E. coli* strains with carbon transporter knockouts

To test whether these simulation predictions can be observed experimentally, we performed controlled *E. coli* co-culture assays. Specifically, we tested how deletion of glucose- and succinate-transporter genes affects niche differences, fitness differences, and coexistence outcomes by running reciprocal invasion assays on three strain combinations under equal (27.5 mM : 27.5 mM) and/or skewed (5 mM glucose : 50 mM succinate) resource ratios (Figure 5A and Figure S2). The first strain combination (Pair #1, denoted by a circle) consisted of two wild-type (WT) strains, each containing all four carbon transporters, and distinguished only by one strain carrying a neutral selection marker. Thus, this strain pair represents the case of neutrality. The second combination (Pair #2, denoted by a square) included one deletion strain lacking a glucose transporter, Δ*manX*, and another lacking a succinate transporter, Δ*dauA*. This pair had a small niche difference due to small negative frequency dependence in the reciprocal invasion assay (Figure S2; top right panel). Unexpectedly, monoculture growth assays revealed that Δ*manX* did not behave as a simple glucose uptake-impaired mutant. Instead, Δ*manX* grew faster than WT in both glucose and succinate, indicating a pleiotropic shift in carbon-use metabolism that is not fully represented by the parameterization of transporter-uptake in COMETS (Figures S3-4; Supplementary Text B). The third combination (Pair #3, denoted by a diamond or an X) consisted of one deletion strain lacking a more dominant glucose transporter, Δ*ptsG*, and the other a different succinate transporter, Δ*dctA*. Strain pair #3 has a large niche difference as both strains grew much more slowly in monoculture on the carbon source matching their deleted transporter. All three pairs were assayed in equal resource ratios, and pair #3 was also assayed in a skewed ratio (denoted by an X).

**Figure 5.**
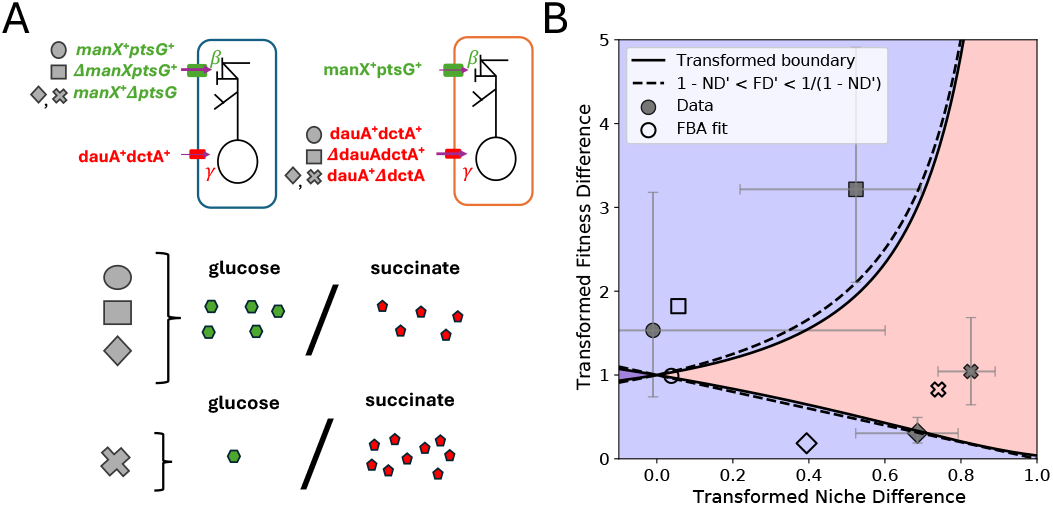
Experimental results from batch co-cultures of *E. coli* strains with targeted carbon transporter knockouts on glucose and succinate. **(A)** Schematic of the experimental conditions, including strain pair genotypes and resource concentration ratios. Gray circle denotes neutral strains where genes encoding all transporters (e.g. *manX, ptsG, dauA* or *dctA*) are present. Gray square corresponds to a small niche difference case where *manX* was knocked out of strain 1 and *dauA* was knocked out of strain 2. Gray diamond and X correspond to a large niche difference case where *ptsG* was knocked out of strain 1 and *dctA* was knocked out of strain 2. All strain pairs were assayed with equal resource concentration ratio (◯, □, ◊), where glucose and succinate were both at 27.5 mM. The third strain pair was also assayed with a skewed resource concentration ratio (X), where glucose concentration was 5 mM and succinate concentration was 50 mM. **(B)** Coexistence phase diagram where gray filled points with 95% confidence intervals correspond to experimental data and open points correspond to dFBA simulation predictions calculated from monoculture data on a single resource to estimate *T*_glucose_ and *T*_succinate_ for each knockout strain.

For each strain pair and resource condition, we used the reciprocal invasion assays to estimate niche differences, fitness differences, and coexistence outcomes in the same transformed phase space used for the dFBA simulations (Figure 1D). We counted colony forming units (CFUs) on selective and non-selective plates to calculate relative initial and final frequencies of each strain which we then used to estimate niche differences, fitness differences and coexistence metrics (see Methods for details). We plotted the outcome of each competition experiment (filled gray markers) in our transformed phase space in Figure 5B, together with dFBA output (open markers) for invasion assays parametrized by monoculture data (Supplemental Text B and Figures S3-4; see Figure S5 for the raw ND-FD phase space). Consistent with the dFBA predictions, we observed the same limiting similarity effect in the experiment, where small and intermediate niche differences (pairs #2 and #3) under equal resource concentrations lead to exclusion. Specifically in pair #2, we observed the exclusion of Δ*dauA* rather than Δ*manX*. The faster growth of Δ*manX* on both glucose and succinate enabled it to outcompete Δ*dauA*. The larger niche and fitness differences of the experimental data compared to the dFBA prediction here can be explained by the fact that no dFBA fit could produce a growth rate as fast as the experimentally-observed growth rate of the Δ*manX* strain on succinate, 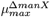 (Figure S4; middle right panel). The large niche difference (pair #3) in skewed resource ratio (X marker) led to coexistence as the high relative concentration of succinate equalized fitness differences between strains. Overall, these findings illustrate that by tuning transporter-mediated trait differences alongside resource concentration ratios, we can predictably modulate niche differentiation and engineer coexistence outcomes in microbial communities.

## Discussion

We developed and validated a mechanistic batch-culture coexistence framework that accounts for microbial feast-famine dynamics and predicts the emergence of limiting similarity from genome-scale metabolic constraints. We show how limiting similarity results from resource quality (maximal growth rates across carbon sources) constraining the response trajectory of interspecific fitness and niche differences to trait changes. The shape of these trajectories can force small trait differences into a region of competitive exclusion and explain the emergence of a minimum niche difference for coexistence that increases with asymmetry in resource quality. Unlike equilibrium theory, coexistence outcomes under fast-famine dynamics depend on initial biomass and resource supply because these conditions determine the duration of transient growth, which amplifies fitness differences relative to niche stabilization. Finally, experimental competition assays with *E. coli* validate our theoretical prediction that metabolic trait perturbations and resource ratios jointly shape trajectories through coexistence space. Taken together, these findings highlight how integrating theory and experiment can reveal the mechanistic metabolic links between trait and resource dynamics, offering a general framework for understanding biodiversity maintenance in complex, nonequilibrium ecosystems.

### Limiting similarity as a mechanistic coexistence criterion

Limiting similarity is generally defined phenomenologically as a minimum distance in trait space [1, 13, 29] or as a minimum difference in resource demands and impacts at equilibrium [2, 30] between species that allow for stable coexistence. In our study, limiting similarity emerges from the joint response of niche and fitness differences to traits and the environment. We show how the nonlinear shape of trajectories leaving neutrality can impose a limiting similarity by preventing small trait changes from directly leading to coexistence. This limiting similarity emerges from trait-based metabolic pathways, mapping genotype-level metabolic differences to phenotypic and ecological responses. In our system, the minimum niche differentiation needed for coexistence scales with how unequal the resources are in supporting growth. When resources differ strongly in quality, greater niche separation is required to compensate for the resulting fitness asymmetry.

The emergence of limiting similarity from nonlinear trajectories in niche–fitness space reframes classic debates about neutrality and niche differentiation [8, 7]. In modern coexistence theory, neutrality represents the limiting case in which niche and fitness differences vanish, suggesting that infinitesimal trait divergence could smoothly generate coexistence. However, when stabilizing and equalizing mechanisms are interdependent, departures from neutrality need not follow orthogonal axes in coexistence space [7]. Our results extend this perspective and apply it to nonequilibrium conditions (e.g. feast-famine dynamics) by showing that the exclusion region is shaped by intrinsic differences in resource quality. Trait perturbations initially amplify fitness asymmetries determined by maximal growth-rate differences between resources, which reflect their underlying chemical properties. In this view, limiting similarity is not imposed as a fixed trait distance but emerges dynamically from how resource chemistry structures the coupling between niche and fitness differences.

### Transient dynamics and initial conditions shape coexistence under feast–famine dynamics

We showed how nonequilibrium feast-famine dynamics constrain the relationship between species’ traits and a community’s trajectory in fitness-niche difference space, which defines the conditions for coexistence. Since feast-famine dynamics are prevalent in natural microbial habitats [21, 20, 19] and are also used in most laboratory experiments [31, 24, 32, 33, 34, 35], the constraints on the coexistence phase diagram identified here provide a generalizable tool to predict coexistence under more realistic conditions found in both natural and experimental communities. Although much work has extended coexistence criteria to nonequilibrium systems [36, 37], an interesting outcome from our study is that the initial conditions (biomass and resource concentrations) alter coexistence predictions. Indeed, they do so by modulating the duration of the transient growth phase, which in turn differentially shapes niche and fitness differences. This is compatible with the perspective that transient dynamics resulting from initial conditions are important in structuring ecological communities [38, 39, 40].

Future work that explicitly manipulates the duration of the exponential growth phase would provide a direct experimental test of our predictions. A recent study varied dilution factors in a two-species, single-resource system [31]. By lowering initial biomass, they effectively extended the exponential growth phase and found that higher dilution factors favoured the species with the highest maximal growth rate, consistent with an amplification of fitness differences during prolonged growth. In contrast, at higher initial biomass (shorter growth phases), they observed coexistence, and in some cases even competitive dominance of the slower-growing species, suggesting the presence of an underlying trade-off that counteracts pure growth-rate advantages.

Extending such experiments to environments with multiple limiting resources, while explicitly quantifying niche and fitness differences through invasion assays, would allow a direct test of how transient growth dynamics mediate coexistence in feast–famine conditions. In our experiments, we used relatively modest dilutions, such that population sizes remained above the range where stochastic processes such as demographic drift become relevant. However, follow-up experiments starting from smaller initial population sizes, which are expected to be more sensitive to drift, could add nuance to our predictions where the effects of stochasticity may limit similarity in a more restrictive way, as shown in recent theoretical work [41]. Furthermore, COMETS [28], the dFBA framework we used, can add drift to the simulations, which would provide a benchmark for experimental studies exploring this question. More broadly, such future extensions of this work would clarify how deterministic metabolic asymmetries and stochastic demographic processes jointly structure coexistence under realistic, nonequilibrium ecological conditions.

### Engineering coexistence through targeted genetic edits

By introducing targeted transporter knockouts and systematically varying glucose-succinate ratios, we engineered predictable coexistence-exclusion outcomes that mapped onto predicted trajectories in niche–fitness space. Although our empirical tests focused on a single carbon source pair, the observed coexistence thresholds were generally consistent with model predictions. Additional high-throughput experiments across varying resource pairs will be necessary to confirm the generality of the relationship between resource quality asymmetry and degree of limiting similarity found in our simulations. Moreover, the Δ*manX* strain revealed an important distinction between knockout expectations, fitted dFBA predictions, and experimentally observed pleiotropy. Although deletion of a glucose transporter was expected to reduce performance on glucose, Δ*manX* instead grew faster than WT on both glucose and succinate. The fitted dFBA simulations captured part of this phenotype: Δ*manX* was assigned a higher glucose uptake bound than WT, which allowed dFBA to correctly predict that Δ*manX* would outcompete Δ*dauA* in batch culture. However, dFBA could not reproduce the experimentally observed gain in succinate growth because both WT and Δ*manX* saturated at the imposed maximum uptake bound, *T* = 10. Thus, Δ*manX* did not fall entirely outside the dFBA framework; rather, its glucose advantage could be absorbed by fitted uptake bounds, whereas its enhanced succinate performance reflected a pleiotropic physiological shift beyond what could be represented by changing uptake alone. Importantly, this also shows that genetic perturbations can tune resource-use traits upward as well as downward. When pleiotropic effects were minimal, experimental outcomes matched dFBA predictions more directly, suggesting that deviations between genotype-phenotype expectations and observed outcomes arise when engineered genotypes alter physiology beyond specific carbon-source uptake. Nonetheless, these deviations highlight opportunities to integrate metabolic pleiotropy with mechanistically explicit genotype-phenotype mappings into coexistence models, which could refine predictions by bridging levels of biological organization into a cohesive framework. In the present system, cross-feeding was negligible, allowing us to interpret coexistence outcomes primarily through direct competition for supplied resources (Figure S6). In communities where metabolic byproducts strongly mediate interactions, future extensions could expand our framework to quantify how secondary resource production reshapes trajectories through ND–FD space by introducing facilitative effects alongside competition [42]. Experimentally, targeted genetic perturbations offer a scalable way to tune competitive trait differences among strains[43, 44], while metabolic gene loss can be used to generate obligate cross-feeding interactions [45]. Together, these approaches suggest a route to designing microbial communities with controlled competitive outcomes.

Overall, our study shows that coexistence in nonequilibrium systems can be mechanistically grounded in metabolic constraints. Rather than treating limiting similarity as an abstract trait distance, we demonstrate that it emerges from chemically determined growth asymmetries and their interaction with transient ecological dynamics. By linking intracellular resource use to trajectories in niche–fitness space and validating these predictions experimentally, we bridge genome-scale metabolism and modern coexistence theory. This integration provides not only a framework for engineering stable microbial communities, but also a general lens for understanding how metabolic structure shapes biodiversity maintenance under realistic, fluctuating environmental conditions.

## Methods

### Conceptual Overview

MCT predicts species coexistence based on how each species limits its own growth rate compared to that of its competitor, which can be partitioned into niche differences (*ND*) and fitness differences (*FD*). MCT is often parameterized using the two-species Lotka–Volterra (LV) competition model, in which

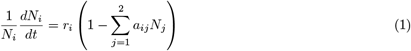

where *N*_*i*_ is the biomass of species *i, r*_*i*_ is its intrinsic growth rate, and *a*_*ij*_ is the relative reduction in species *i* ‘s intrinsic growth caused by one unit of density of species *j*. From this model, *ND* and *FD* are respectively defined as

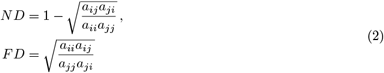

and stable coexistence is predicted when

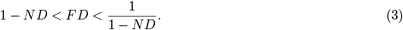

Since competition coefficients *a*_*ij*_ and *a*_*ji*_ are phenomenological, any combination of *ND* and *FD* is, in principle, achievable. Therefore, the coexistence space is unconstrained.

Mechanistic consumer-resource theory, however, suggests that a single trait change often alters both negative frequency dependence, which corresponds to *ND*, and relative competitive ability, which corresponds to *FD*. Here, we focus on a community of two species and two carbon sources, *β* and *γ*, where *T* is the uptake rate of an alternative (less preferred) carbon source *β*. We define *T*_*max*_ as the maximum uptake rate on the species’ preferred carbon source (if *β* is less preferred then uptake rate of *γ* is *T*_*max*_). Assuming that carbon is the limiting element, changes in the uptake trait *T* and in the concentration ratio of the two carbon sources 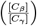 channel species away from neutrality along specific trajectories. We write this constraint schematically as:

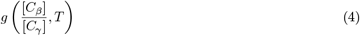

implying that departures from neutrality are not free to roam the niche-fitness space independently (Eq. 3). To map *g*, we combine (i) genome-scale metabolic modeling and (ii) controlled invasion experiments, which we detail in the following sections.

### Dynamic Flux Balance Analysis simulations

We used the *E. coli* genome-scale reconstruction iJO1366 as the base metabolic model and embedded it in COMETS v0.12 for all dFBA [28]. Simulations were executed through the Python API (`cometspy`) with the Gurobi 10.0.3 solver, maximizing biomass flux at each 0.1h time step. Each batch run spanned 24 h (240 time steps), capturing a single feast–famine cycle.

#### Trait manipulation

COMETS represents resource uptake as a lower bound (negative flux indicates import). For readability, we report the magnitude of this bound as a positive rate. In each simulated competition, we created two otherwise isogenic strains: in strain 1, the maximum uptake magnitude on carbon source *β* was reduced to a value *T* (varied from 10 to 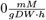), while strain 2 carried the same reduction on carbon source *γ*. The units of flux correspond to millimoles of metabolite consumed per gram of biomass per hour. All other exchange reactions retained an unconstrained uptake magnitude of 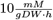. We say ‘unconstrained’ here because above a flux value of 10, the resource import reaction is no longer limiting for growth; rather, some other bottleneck in the network becomes limiting [28]. Thirty-six carbon source pairs were examined (all unordered pairwise combinations of the nine carbon sources from Figure 3 are listed in Table S1).

#### Resource environments

For each pair of resources, we imposed 11 evenly spaced resource concentration ratios spanning [*C*_*β*_] : [*C*_*γ*_] = 1 : 10 to 10 : 1 while maintaining the total added carbon constant at 55 *mM*. Media trace ions and vitamins were supplied in excess (1 M and non-depleting) to ensure carbon limitation. The initial total biomass was 5 *×* 10^−6^ gDW, partitioned to produce reciprocal invasion frequencies of 1:99 and 99:1 (robustness to the initial frequency shown in Figure S7). COMETS logged species biomasses, extracellular metabolite concentrations, and fluxes at each simulation cycle. Code is available at https://github.com/brendonmcguinness/coexistence-dFBA-exp.

#### In silico invasion metrics: selection coefficients, raw niche & fitness differences

Reciprocal invasion assays were produced *in silico* by performing two batch simulations, *k* ∈ *{L, H}*, per strain pair and resource ratio: one for the low-frequency invader case, *k* = *L*, with initial frequency *p*^(*L*)^ = 0.01 of total initial biomass, and one for the high-frequency resident case, *k* = *H*, with *p*^(*H*)^ = 0.99 of total initial biomass. Let 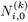 and 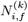 denote the initial (*t* = 0) and final (*t* = *f*) biomasses (gDW) of strain *i* over the simulated batch assay under frequency case *k*. The Malthusian growth of strain *i* under frequency *k* is

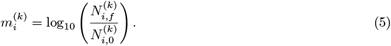

The selection coefficient of strain *i* relative to strain *j* in that invasion assay is

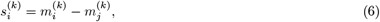

so 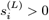 indicates that *i* increases in proportion when introduced at low frequency (*k* = *L*) into a population dominated by *j*. From the two reciprocal invasion assays, we obtain 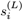 when strain *i* is rare and 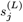 when strain *j* is rare. Note that 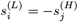 because when *k* = *L* for strain *i, k* = *H* for strain *j*, which experiences the same selective pressure but in the opposite direction.

Negative frequency dependence is determined from the change in selection coefficient across the two initial frequencies. Using the initial frequencies as inputs for each simulation, *p*^(*L*)^ and *p*^(*H*)^, we define the raw niche difference, *ND*, as the negative frequency-dependence slope:

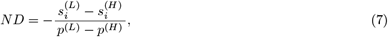

where selection for strain *i* improves as its initial frequency declines (stabilizing). The raw fitness difference, *FD*, is the competitive advantage of strain *i* across both invasion contexts:

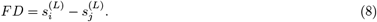

Finally, we define a coexistence strength as

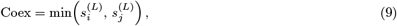

such that Coex *>* 0 indicates mutual invasibility and predicted coexistence, while Coex *<* 0 indicates exclusion of one strain when rare. Figure 2 from the Main text is shown on these raw axes in the SI as Figure S8.

### Mapping dFBA to the Lotka-Volterra *ND***-***FD* **coexistence space**

Selection coefficients are log_10_ fold-changes in biomass bounded by resource supply; as trait difference increases, the invader progressively exploits its preferred resource, driving 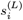 to a ceiling set by resource stoichiometry. Consequently, *ND*, which is proportional to the difference 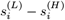, saturates with trait difference rather than growing without bound. We therefore normalize using a saturating function anchored to the maximum observed value *ND*_max_,

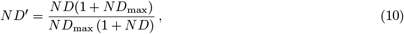

so that *ND*^*′*^ → 1 as niche overlap vanishes and *ND*^*′*^ → 0 as niche overlap is complete. Raw fitness differences are exponentiated to yield a positive scale comparable to LV fitness ratios,

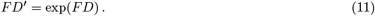

These transformed *ND*^*′*^ and *FD*^*′*^ values are used throughout the Results (unless otherwise noted) to plot mechanistic trajectories in coexistence phase space and to compare directly with the LV coexistence inequality (Eq. 3).

### Experimental strain construction

All strains were derived from *E. coli* MG1655 (parental background; Table S2). We generated single, in-frame deletions of four carbon transporter genes: *manX* (glucose phosphotransferase system), *dauA* (succinate transporter), *ptsG* (high-affinity glucose transporter), and *dctA* (C4-dicarboxylate transporter) using the *λ*-Red system [43]. Briefly, PCR-amplified antibiotic resistance cassettes flanked by 50-bp homology arms were electroporated into arabinose-induced *recA*^+^ cells carrying the temperature-sensitive helper plasmid pKD46; recombinants were selected on plates with kanamycin. Where noted, the resistance cassette (kan^*R*^) was excised by FLP recombinase (expressed from plasmid pCP20), leaving a single FRT scar. In several strains, the resistance cassette was maintained to differentiate strains in co-culture experiments.

To enable reciprocal invasion assays in the wild-type strain, we introduced a neutral selectable marker (kan^*R*^) into the alternate strain (Table S2). Colony PCR across the deletion junctions (see Table S3 for primers used) confirmed correct deletions in all experimental strains. Representative gel is shown in Figure S9. Strains were stored at −80°C in 25% glycerol and streaked before for each experimental block to minimize laboratory evolution.

### Growth assays (monoculture)

We phenotypically validated all transporter knockouts in carbon-defined monocultures using 96-well plate growth assays. The strains were streaked from frozen stocks and a single colony was grown overnight in LB medium, then diluted 1000*x* into EZ rich defined medium minus amino acids (EZRDM^−*AA*^) with either glucose or succinate at 27.5 mM. After 24 h, 5 *µL* of each strain at 1 : 10 dilution was inoculated into wells containing 195 *µL* EZRDM^−*AA*^ (overall dilution 1 : 400) supplemented with a single focal carbon source at 27.5 mM (the concentration used in later co-culture assays). 96-well plates were incubated at 37°C with orbital shaking at 840 RPM in a plate reader with 1mm orbital diameter on the double orbital setting (BioTek Synergy H1) and OD_600_ was recorded every 15 min for 30 h. Blank wells (medium only) were included by column and subtracted from all readings after pathlength normalization. Outer wells on the plate were not used to minimize variability caused by uneven evaporation across the plate (6 technical wells per strain per resource).

To compare experimental data with dFBA simulations, OD_600_ was converted to biomass using empirically determined conversion factors specific to the growth substrate (*a*_glc_ = 0.005 gDW/OD; *a*_suc_ = 0.002 gDW/OD). A hierarchical coarse-to-fine grid search was then used to identify the maximum uptake flux bound *T* in COMETS that minimized the squared difference between simulated and observed final biomass for each strain. Best-fit *T* values and associated errors are reported in Table S4. Each transporter knockout showed a reduction in maximum growth rate, *µ*_max_, on its target resource relative to wildtype, with limited pleiotropic effects (except for Δ*manX*, as discussed in the main text) on the alternate resource (Figures S3-4). These experimentally quantified trait differences define the *T* values used to parameterize our dFBA simulations and to test constrained departures from neutrality in co-culture competition experiments.

### Reciprocal invasion experiments (co-culture)

We quantified pairwise coexistence empirically by running single feast-famine batch co-cultures in defined carbon media with reciprocal starting frequencies: strain 1 invader (5%) versus strain 1 resident (95%), similar to the *in silico* design but not as drastic to ensure enough colony forming units (CFU) on each plate. All assays used the EZRDM^−*AA*^ base medium supplemented with either equal resource concentrations (27.5 mM each) or a skewed glucose:succinate mix (5 mM:50 mM; total 55 mM carbon). Antibiotics were omitted during competition; kan^*R*^ markers were used only to differentiate strains on selective plates after competition. Frozen stocks were streaked, single colonies grown overnight in carbon-replete pre-culture medium, washed twice in carbon-free EZRDM base (4,000 *g*, 4 min, room temperature), and resuspended to OD_600_ = 0.2 then diluted so that starting OD_600_ = 0.02. Cell densities in co-culture were measured by CFU plating. For each strain pair, we prepared two reciprocal mixes targeting 5:95 and 95:5 (invader:resident) in 1.5 mL eppendorf tubes. Cultures were incubated at 37°C with orbital shaking (840 RPM) for 30 h. At *t*_0_=0 h and *t*_*f*_ =30 h, aliquots were serially diluted in 0.085% saline and plated (*n* = 6) on: (i) non-selective LB (total CFU) and (ii) selective LB+kan. Plates were incubated for 20h at 37°C and CFUs were counted by automated imaging with manual curation (Supplemental Text C and Figure S10).

#### Coexistence metrics from plate counts

Initial 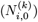 and final 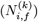 biomass for each strain were calculated from CFU counts corrected for dilution factors and plating volumes. We then computed Malthusian growth *m*_*i*_ and selection coefficients 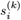 exactly as defined for *in silico* invasions (Eqs. 5, 6), simply substituting CFU for biomass. Raw niche and fitness differences (*ND, FD*) and coexistence strength (Coex) were then obtained from Eqs. 7-9 using these experimental values. Transformed LV-scale *ND*^*′*^ and *FD*^*′*^ were computed with Eqs. 10-11 for direct comparison to the theoretical coexistence boundary. Each strain pair *×* resource concentration ratio *×* initial frequency was assayed in 2 biological replicates each with three technical replicates on separate days, *n* = 6 (with the exception of the neutral strains which were assayed in three technical replicates on one day, *n* = 3). We calculated 95% confidence intervals for each data point.

## Data, Materials and Software Availability

All code and data used to generate figures and conduct analysis are available at https://github.com/brendonmcguinness/coexistence-dFBA-exp/.

## Acknowledgments

We thank members of the Guichard and Weber labs at McGill University for helpful discussion. We graciously thank Dr. Fiona Soper for allowing us to use her lab’s plate reader for the experiments conducted in this study. This work was supported by the Natural Sciences and Engineering Research Council of Canada RGPIN-2024-05623 to S.C.W. and RGPIN-2023-03622 to F.G. This research was undertaken, in part, thanks to funding from the Canada Research Chairs Program to S.C.W. We also thank Quebec Oceans for their financial support.

## Author Contributions

B.M., F.G., and S.C.W. designed the research. B.M. performed the research and drafted the paper. S.C.W. trained B.M. on the experimental tools to perform the research. B.M., F.G., and S.C.W. contributed to writing and revising the paper.

