## Supplemental Text for "A winding road to coexistence: Interdependence of niche and fitness differences in *E. coli* with targeted resource uptake gene deletions"

#### Supplementary Text A: Predictors of maximal growth rate from cofactor, substrate, and network features

The dFBA solution returns a flux vector over thousands of reactions in iJO1366, from which growth rate emerges as an output of the linear program [1]. Although this high-dimensional flux vector specifies the metabolic state, it is not immediately clear why one carbon source supports faster growth than another, or which coarse features of the resource and flux solution are most strongly associated with these differences. We therefore asked which biologically interpretable descriptors of cofactor production, substrate chemistry, and metabolic network organization covary with maximum growth rate across carbon sources. This analysis provides a bridge between genome-scale flux organization and the growth-rate differences that contribute to niche and fitness differences in our coexistence framework.

For each resource $\alpha$ in the extended set of 23 carbon sources (Table S1), which includes the nine carbon sources used in Fig. 3 plus 14 additional carbon sources to increase the number of observations, we ran a one-cycle COMETS simulation [1] using the iJO1366 metabolic model. Simulations used a time step of $\Delta t = 1$ h, an uptake lower bound of $-10$ mM gDW$^{-1}$ h$^{-1}$, initial biomass $N_0 = 10^{-6}$ gDW, and an initial resource concentration of 27.5 mM. Maximum growth rate on carbon source $\alpha$ was approximated from total biomass over this one-hour interval as

$$\mu_{\max}(\alpha) = \log(N_1/N_0), \tag{1}$$

where $N_1$ is the biomass after $\Delta t = 1$ h. Because the initial biomass is small, substrate depletion over this interval is negligible, allowing this one-cycle estimate to approximate the substrate-specific maximum growth rate. We refer to $\mu_{\max}(\alpha)$ as $\mu$ throughout this section.

We first extracted cofactor production features from the flux solution. For each cofactor $Y \in \{\text{ATP}, \text{NADH}, \text{NADPH}\}$, we summed the flux through reactions with a positive stoichiometric coefficient for the cytosolic form of $Y$, giving a gross cofactor production flux $Y_{\text{prod}}$. Because intracellular cofactors are mass-balanced at steady state in the dFBA model, these quantities do not measure net accumulation or excess production. Instead, they measure cofactor turnover: the total pathway flux generating each cofactor that must be balanced by downstream consumption. Therefore, these descriptors summarize how strongly each substrate participates in energy, redox, and biosynthetic cofactor metabolism.

Next, we included substrate-level descriptors. For each substrate, we recorded the number of carbon atoms, #C, as a simple measure of substrate size. We also computed a coarse carbon-efficiency metric,

$$C_{\text{eff}}(\alpha) = \frac{C_{\text{in}} - C_{\text{CO}_2,\text{out}}}{C_{\text{in}}}, \tag{2}$$

where

$$C_{\text{in}} = \#\text{C}(\alpha)\,|v_{\text{EX}_\alpha}|. \tag{3}$$

Here, $C_{\text{CO}_2,\text{out}}$ was estimated from the $CO_2$ exchange flux, treating $CO_2$ as the dominant measured carbon-loss term. Together, #C and $C_{\text{eff}}$ capture coarse aspects of substrate chemistry: how much carbon enters the system and what fraction of that carbon is retained as biomass rather than lost as $CO_2$.

Finally, we computed descriptors of metabolic network organization from the flux log of each single-cycle simulation. We defined the set of active reactions as

$$\mathcal{R} = \{r : |v_r| > 10^{-12}\}, \tag{4}$$

where $v_r$ is the reaction rate (in $\frac{\text{mmol}}{\text{gDW}\cdot\text{h}}$), and recorded the number of active reactions as a measure of the breadth of metabolic network engagement on each substrate. We also calculated total absolute flux,

$$V_{\text{tot}} = \sum_{r \in \mathcal{R}} |v_r|, \tag{5}$$

and flux entropy,

$$H_v = -\sum_{r \in \mathcal{R}} p_r \log p_r, \qquad p_r = \frac{|v_r|}{\sum_{r' \in \mathcal{R}} |v_{r'}|}. \tag{6}$$

Flux entropy measures how evenly total flux is distributed across active reactions. Low entropy indicates that flux is concentrated through a small number of dominant reactions, whereas high entropy indicates a more distributed metabolic state. Thus, active reaction number and flux entropy capture global properties of flux organization that are not reducible to cofactor production alone. They allow us to distinguish substrates processed through focused, high-throughput routes from those requiring broader redistribution of flux across the network.

To further quantify pathway structure, we built an undirected reaction graph with active reactions as nodes and edges between reactions that share a metabolite. Edge weights were defined as

$$w(r \to r') = \frac{1}{|v_{r'}|}, \tag{7}$$

so that high-flux routes correspond to shorter effective paths. We then computed a weighted shortest-path length $d_s$ from the uptake reaction $\text{EX}_\alpha$ to the biomass reaction using Dijkstra's algorithm [2]. We also calculated a bottleneck flux, defined as the minimum $|v_r|$ along this shortest path, excluding the uptake and biomass reactions. These quantities summarize, respectively, an effective pathway resistance from resource uptake to biomass and the tightest internal flux constraint along that path.

The resulting per-substrate data frame included $\mu$ and the following candidate predictors: ATP production, NADH production, NADPH production, #C, $C_{\text{eff}}$, shortest-path length $d_s$, bottleneck flux, total flux, flux entropy, and number of active reactions.

Because the number of carbon sources ($N = 23$) was small relative to the number of candidate descriptors, we used simple, interpretable regression analyses rather than high-capacity predictive models. We first assessed each descriptor individually by fitting a linear regression of $\mu$ against that predictor and recording the Pearson correlation coefficient, in-sample $R^2$, RMSE, and leave-one-out cross-validated prediction error. This analysis identifies which individual cofactor, substrate, or network descriptors are most strongly associated with variation in maximum growth rate, without requiring a multivariate model that would be difficult to estimate robustly from the available number of substrates.

Among the descriptors considered, NADH production showed the strongest positive association with $\mu$ (Fig. S1A–B; Table S5), explaining 61% of the observed variation in maximum growth rate in-sample. This relationship retained predictive value under leave-one-out cross-validation, with $R^2_{\mathrm{LOOCV}} = 0.47$. Carbon efficiency showed a similar single-predictor association, but, as described below, added less information once NADH production was already included in the model.

Next, we ask whether any substrate- or network-level descriptor explained residual variation beyond NADH production. To do so, we fit two-predictor models of the form

$$\mu \sim \mathrm{NADH}_{\mathrm{prod}} + x_j, \tag{8}$$

where $x_j$ was each remaining descriptor in turn. We compared each two-predictor model to the NADH-only model using in-sample $R^2$, adjusted $R^2$, Akaike's information criterion corrected for small sample size (AICc), and leave-one-out cross-validated prediction error. Leave-one-out cross-validation was used to determine whether an added descriptor improved prediction of held-out substrates, rather than merely improving fit to the observed substrate panel.

Adding the number of active reactions produced the strongest improvement beyond NADH production (Table S5). The model $\mu \sim \mathrm{NADH}_{\mathrm{prod}} + \#\text{active reactions}$ increased in-sample $R^2$ from 0.61 to 0.70 and $R^2_{\mathrm{LOOCV}}$ from 0.47 to 0.59, while reducing LOOCV RMSE from 0.132 to 0.115 and reducing AICc by approximately 3 units. Carbon efficiency produced a smaller improvement, whereas the remaining descriptors did not improve held-out prediction relative to the NADH-only model. Thus, active reaction number captured residual network-level information about growth rate beyond redox production alone.

We also performed principal component analysis (PCA) on the standardized predictor matrix as an unsupervised complement to the regression analyses. PCA was used to describe covariance among cofactor, substrate, and network descriptors; $\mu$ was not included in the PCA. PC1 explained 32.5% of the variance among metabolic descriptors and loaded positively on NADH production, carbon efficiency, carbon number, and active reaction number (Fig. S1C), indicating that these features covary along a broad substrate-efficiency/redox axis. However, PC1 was less predictive of $\mu$ than NADH production alone, showing that the dominant axis of metabolic-feature variation was not identical to the growth-rate axis.

Overall, this analysis asks whether variation in maximum growth rate is primarily associated with specific energetic and redox outputs, coarse substrate chemistry, or broader network-level properties such as total flux, flux entropy, and active reaction number. NADH production was the strongest single descriptor of $\mu$, suggesting that substrate-dependent growth differences are closely tied to redox yield. However, active reaction number improved prediction beyond NADH production, indicating that the breadth of active metabolic network engagement also contributes to growth-rate variation. Notably, although carbon efficiency $C_{\text{eff}}$ showed a strong raw correlation with $\mu$, it produced a smaller added-value improvement once NADH production was already in the model. This pattern is consistent with the PCA result: $C_{\text{eff}}$ and $\text{NADH}_{\text{prod}}$ both load strongly on PC1 and therefore carry partly overlapping information, whereas the number of active reactions captures a network-level component of variation that is not redundant with NADH production. In this way, substrate chemistry propagates through intracellular metabolism to generate growth-rate asymmetries, which in turn contribute to the fitness differences measured in the coexistence analyses.

#### Supplementary Text B: Monoculture fitting of transport bounds

Experimental monoculture time series on one carbon source, either glucose or succinate, were used to fit strain-specific transport bounds for COMETS simulations. The co-culture assays included three strain-pair contexts: Pair #1 was the neutral WT marker-control pair; Pair #2 was the weak transporter-deletion contrast, $\Delta manX$ versus $\Delta dauA$; and Pair #3 was the stronger transporter-deletion contrast, $\Delta ptsG$ versus $\Delta dctA$. These fitted bounds allowed us to parameterize dFBA simulations using experimentally measured monoculture growth phenotypes and then compare the resulting co-culture predictions to the experimental outcomes shown in Figure 5 of the main text. For each strain–resource combination, we used a grid search to identify the maximum uptake rate,

$T$, that minimized the root mean squared error (RMSE) between simulated and observed OD over the growth phase. Unless otherwise noted, the fitting window was defined from $t\_0 = 0.24$ h to the first time point at which OD reached 0.40. This window isolates the exponential portion of growth, where the uptake bound most directly controls the trajectory, while avoiding lag and stationary phases.

The only exception was $\Delta dctA$ on succinate. This growth curve showed a pronounced extended lag, with OD remaining near baseline until approximately 20 h, followed by a brief rise to approximately OD 0.3 (Figure S3, bottom row). Restricting the fit to this narrow late-growth window produced an unrepresentative bound because the simulation, which does not include an explicit lag phase, was forced to fit only the late steep rise while systematically mismatching the early trajectory. We therefore fit $\Delta dctA$ on succinate using the full time series, which produced a bound that better captured its overall growth behavior. This exception did not change the qualitative ordering of fitted bounds across strains, but only affected the $\Delta dctA$ point estimate.

The best-fit values of $T$ were used to parameterize simulations for the corresponding co-culture assays. Fitted values and errors are reported in Table S4. Values of $T > 10.00$ produced the same maximum growth rate in COMETS, so $T = 10.00$ was used as the upper bound of the grid search.

The fitted transport bounds generally had the expected effects of transporter deletions. Strains lacking major transporters for a given carbon source required lower fitted uptake bounds on that resource, consistent with reduced monoculture growth on the corresponding substrate. For example, $\Delta ptsG$ had a reduced fitted glucose uptake bound, whereas $\Delta dctA$ had a reduced fitted succinate uptake bound. Thus, the fitted uptake bounds for Pair #3 showed larger reductions on the corresponding deleted carbon sources than those observed for Pair #2.

The main exception was the $\Delta manX$ strain, which did not behave as a glucose uptake-impaired mutant. Although deletion of a glucose transporter was expected to reduce performance on glucose, $\Delta manX$ grew faster than WT on both glucose and succinate. The fitted dFBA parameterization captured part of this phenotype: on glucose, $\Delta manX$ was assigned a higher fitted uptake bound than WT, allowing the fitted model to represent improved glucose performance for $\Delta manX$ relative to WT. However, on succinate, both WT and $\Delta manX$ reached the imposed maximum uptake bound of $T = 10$. Values of $T > 10$ produced the same maximum simulated growth rate in COMETS, so $T = 10$ was used as the upper limit of the grid search. Consequently, the experimentally observed enhancement of $\Delta manX$ growth on succinate could not be reproduced by further increasing the uptake bound.

Thus, the $\Delta manX$ phenotype was only partially captured by fitting carbon uptake bounds. Its glucose advantage could be absorbed by the fitted uptake parameter, whereas its enhanced succinate growth reflected a pleiotropic physiological shift beyond what could be represented by changing uptake capacity alone. This distinction is important for interpreting Pair #2: rather than representing a clean reciprocal pair of weak glucose and succinate specialists, Pair #2 combines weak niche differentiation with a $\Delta manX$-specific fitness advantage. Therefore, while the theoretical simulations in the main text represent idealized transporter effects through prescribed uptake bounds, the fitted dFBA simulations should be interpreted as an empirical parameterization of monoculture growth phenotypes, rather than as a complete mechanistic explanation of the physiological effects of each deletion.

#### Supplementary Text C: Semi–automated colony counting

Colony forming units (CFUs) at $t_0 = 0$ h and $t_f = 30$ h were counted from digital images (taken with an iPhone 13 camera) of LB agar plates (selective and non-selective). We built a semi–automated ImageJ macro (`https://github.com/brendonmcguinness/coexistence-dFBA-exp`) which combines standardized preprocessing with manual quality control to balance speed and accuracy of colony counting. Briefly, we converted the iPhone .HEIC images into TIFF images and opened them in Fiji. Images were processed by the macro which:

1. Converts to 8-bit grayscale and interactively crops to the plate boundary.
2. Prompts the user to draw an oval region of interest (ROI) enclosing the usable agar surface and clears pixels outside it to remove edge glare.
3. Applies a manually selected foreground threshold and binarizes the image with a white-object mask so that pixels denoting a part of a colony have value 1 and background has value 0. Thresholds were selected interactively per image based on the bimodal intensity histogram, with the operator ensuring separation of colony and background peaks.
4. Performs morphological cleaning so that only colonies that are filled are counted, whereas

those with holes in their center are not counted as they are likely artifacts from bubbles from when the LB media solidified. This is later confirmed during the manual step.

5. Executes a watershed operation to split touching colonies in order to count them as separate colonies.

6. Runs ImageJ's `Analyze Particles` function with empirically chosen filters (`size=30-Infinity` pixels; `circularity=0.50 -1.00`) to detect colonies, adding each particle as an ROI and producing per plate colony count. Detected colonies are overlaid on the original image for visual auditing.

7. Users then conduct a guided quality–control step: false positives are deleted from the ROI Manager; missed colonies are added manually. A second macro `MakePlateSummary.ijm` is used to export a plate–level CSV containing (i) plate identifier, (ii) count, (iii) selective medium type, (iv) dilution factor, and (v) timestamp.

All CFU estimates incorporate the specified dilution and plating volume to yield absolute densities. Only plates with 30–300 counted colonies were used for quantitative analyses, following standard microbiological practice to balance stochastic counting error at low densities and colony merging at high densities. Figure S10 shows an example of a plate and the ROI after step 2 (Figure S10A) and after step 7 (Figure S10B).

#### Supplementary Figures

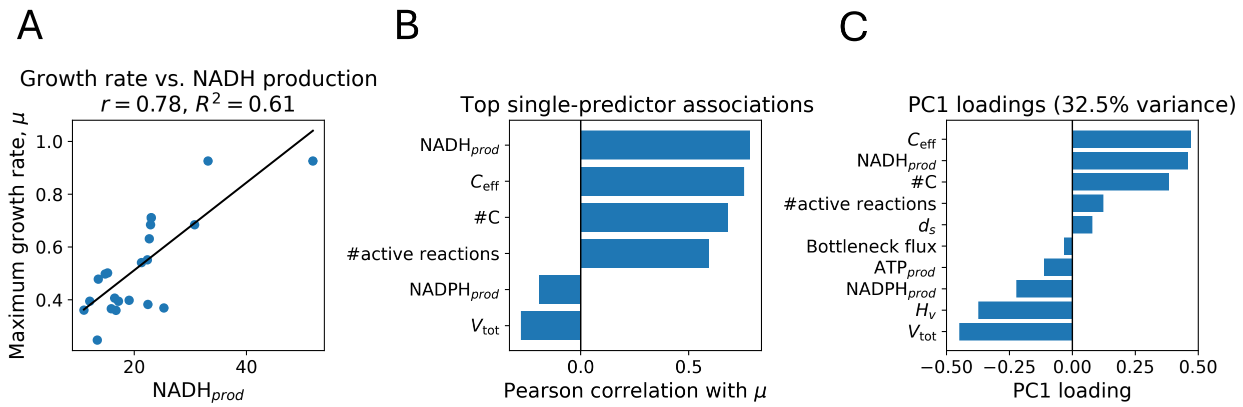


**Figure S1: Cofactor, resource, and network predictors of maximum growth rate across the extended carbon source set.** **(A)** Maximum growth rate $\mu$ increased with gross NADH production across the 23 single-resource dFBA simulations. Points represent individual carbon sources; the line shows the ordinary least-squares fit. **(B)** Pearson correlations, $r$, between $\mu$ and the six strongest single-predictor descriptors. NADH production and carbon efficiency showed the strongest positive single-predictor associations with maximum growth rate, followed by carbon number and active reaction number. **(C)** PC1 loadings from PCA of the standardized predictor matrix. PCA was performed without including $\mu$, so loadings describe covariance among metabolic descriptors rather than growth-rate predictor importance. PC1 loaded positively on NADH production, carbon efficiency, carbon number, and active reaction number, consistent with a broad substrate-efficiency/redox axis.

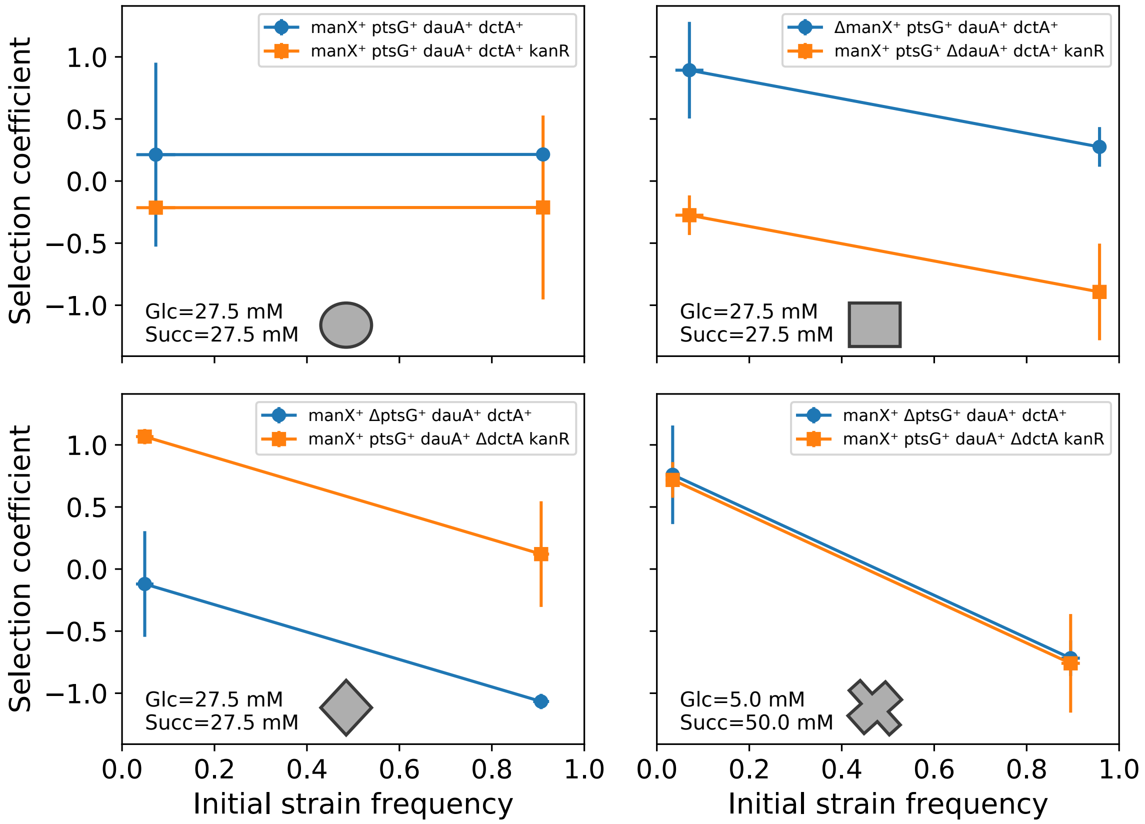


**Figure S2: Reciprocal invasion assays to quantify frequency–dependent selection and coexistence.** Selection coefficient is plotted against initial strain frequency across two resource concentration ratios: equal (Glc = 27.5 mM, Suc = 27.5 mM) or skewed (Glc = 5.0 mM, Suc = 50.0 mM) carbon sources, as labeled. Genotypes atop each panel indicate transporter knockout for the competing strains (*manX*, *ptsG*, *dauA*, dctA, manX, ptsG); the competitor carries a neutral *kanR* marker for plate-based differentiation

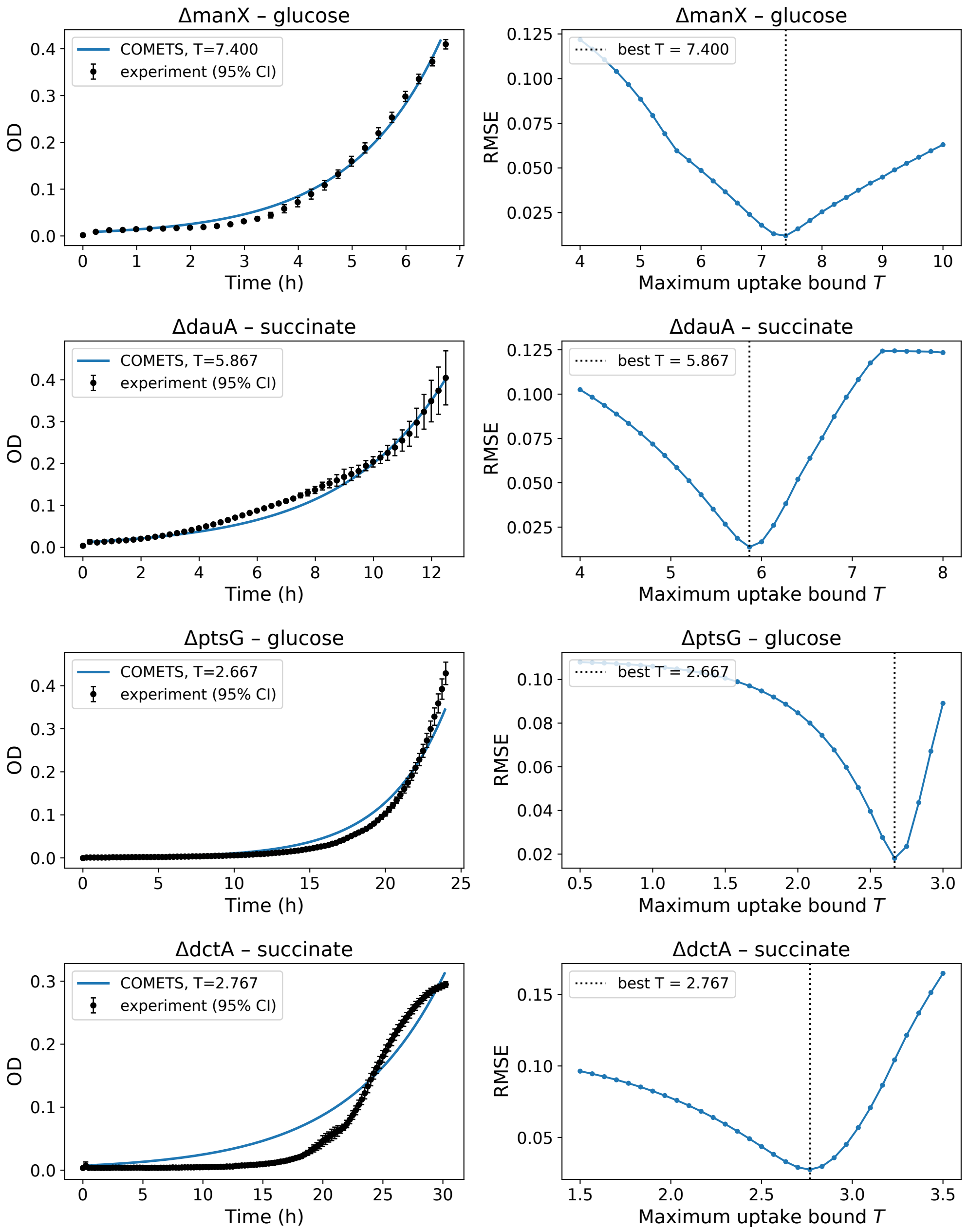


**Figure S3: Fitting of COMETS maximum uptake bounds to monoculture growth curves.** Each row shows one knockout strain grown on the carbon source for which its transporter is impaired (top to bottom: $\Delta manX$ on glucose, $\Delta dauA$ on succinate, $\Delta ptsG$ on glucose, $\Delta dctA$ on succinate). Left column: optical density at 600 nm (OD) versus time (hours). Black points are experimental measurements (mean $\pm$ 95% CI across replicates); the blue curve is the COMETS dynamic-FBA simulation at the best-fit maximum uptake bound T. Simulations were initialized with biomass matched to the experimental OD to fit over the growth phase only with the exception of $\Delta dctA$ which we fit to the whole time series. Right column: root-mean-square error (RMSE) between simulated and observed OD as a function of $T$; the dotted vertical line marks the best-fit value ($\Delta manX$ $T = 7.40$, $\Delta dauA$ $T = 5.87$, $\Delta ptsG$ $T = 2.67$, $\Delta dctA$ $T = 2.77$ mmol gDW$^{-1}h^{-1}$), taken as the RMSE minimum.

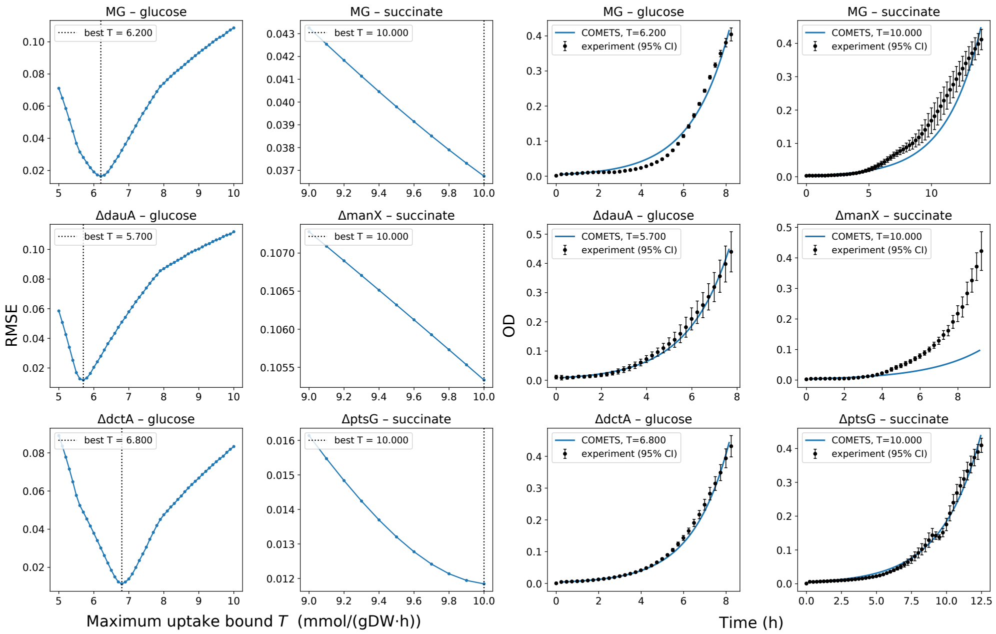


**Figure S4: Fitting maximum uptake bounds on the alternate (intact transporter) carbon source.** Fits for WT on each carbon source and for each knockout grown on the carbon source that it can still transport normally (rows: WT; pair #2, $\Delta dauA/\Delta manX$; and pair #3, $\Delta dctA/\Delta ptsG$). *Left two columns:* RMSE between simulated and observed OD as a function of the maximum uptake bound $T$; the dotted vertical line marks the best-fit value. *Right two columns:* OD versus time, with experimental data (black, mean ± 95% CI) and the COMETS simulation at best-fit $T$ (blue). Simulations were initialized with biomass matched to the experimental OD. On glucose, WT, $\Delta dauA$, and $\Delta dctA$ show well-defined RMSE minima (T = 6.2, 5.7, 6.8 $\frac{\text{mmol}}{\text{gDW}\cdot h}$, respectively). On succinate, WT, $\Delta manX$, and $\Delta ptsG$ show RMSE decreasing monotonically to the search ceiling, so their bounds are reported at $T = 10$ (upper limit) rather than as an interior optimum.

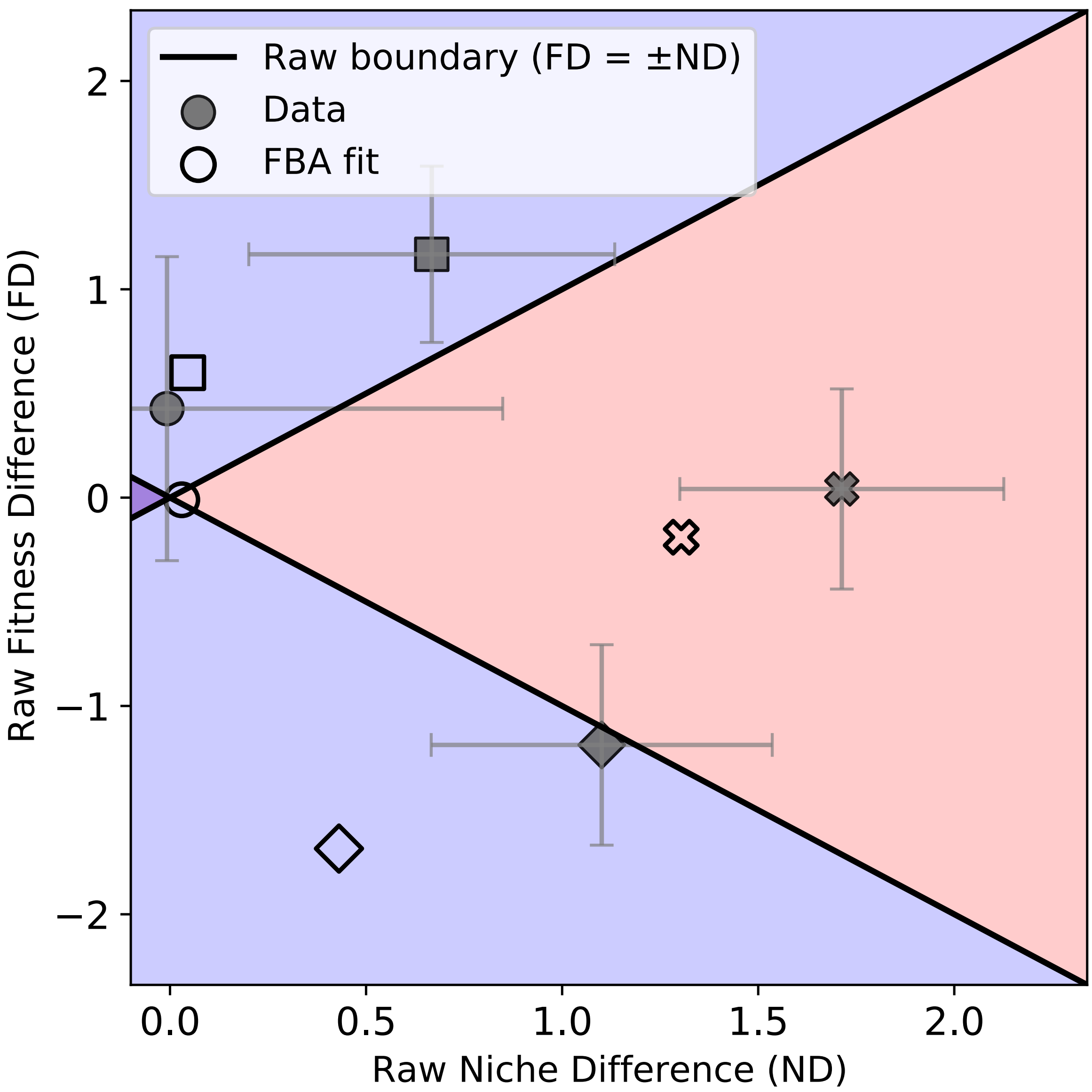


**Figure S5: Raw niche and fitness differences from batch coculture experiments of *E. coli* strains with targeted carbon transporter knockouts on glucose and succinate.** Gray circle denotes neutral strains where all transporter genes (*manX*, *ptsG*, *dauA* or *dctA*) are present. Gray square corresponds to a small niche difference case where *manX* was knocked out of strain 1 and *dauA* was knocked out of strain 2. Gray diamond and gray 'X' correspond to a large niche difference case where *ptsG* was knocked out of strain 1 and *dctA* was knocked out of strain 2. All strain pairs were assayed with equal resource concentration ratio (◯, □, ◊), where glucose and succinate were both supplied at 27.5 mM. The third strain pair was also assayed with a skewed resource concentration ratio ('X'), where glucose was supplied at 5 mM and succinate was supplied at 50 mM. Coexistence phase diagram where gray filled points with 95% confidence intervals correspond to experimental data and open points correspond to dFBA simulation predictions calculated from monoculture data on a single resource to estimate $T_{\text{glucose}}$ and $T_{\text{succinate}}$ for each knockout strain.

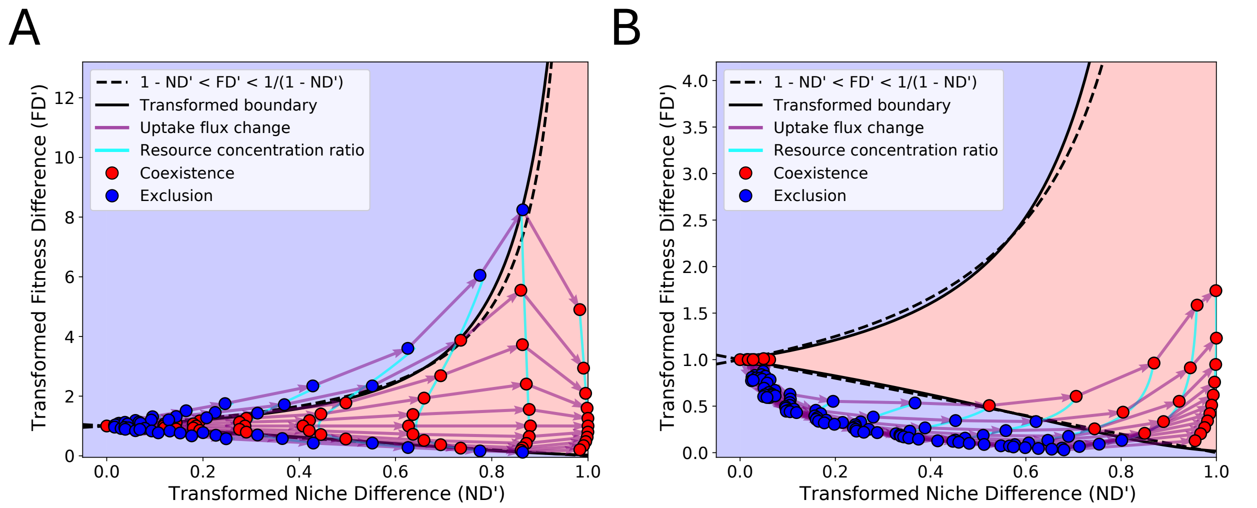


**Figure S6: Departure from neutrality trajectories are qualitatively robust to cross-feeding.** Dynamic FBA trajectories in $ND$–$FD$ space when consumption of secreted by-products is enabled for both strains (i.e. cross-feeding). Shaded regions show the theoretical coexistence (red) and exclusion (blue) regions; dashed curves are the transformed coexistence boundaries. Vectors depict two mechanistic gradients: changing the relative resource concentration ratio at fixed trait (cyan) and decreasing the alternative-resource uptake capacity $T$ at fixed supply (magenta). Markers indicate outcomes from mutual-invasibility tests (red = coexistence, blue = exclusion). **(A)** Resource pair with similar quality (glucose and fructose). **(B)** Resource pair with large maximal-growth asymmetry (glucose and succinate). Allowing cross-feeding slightly shifts magnitudes but does not alter the qualitative geometry of trajectories.

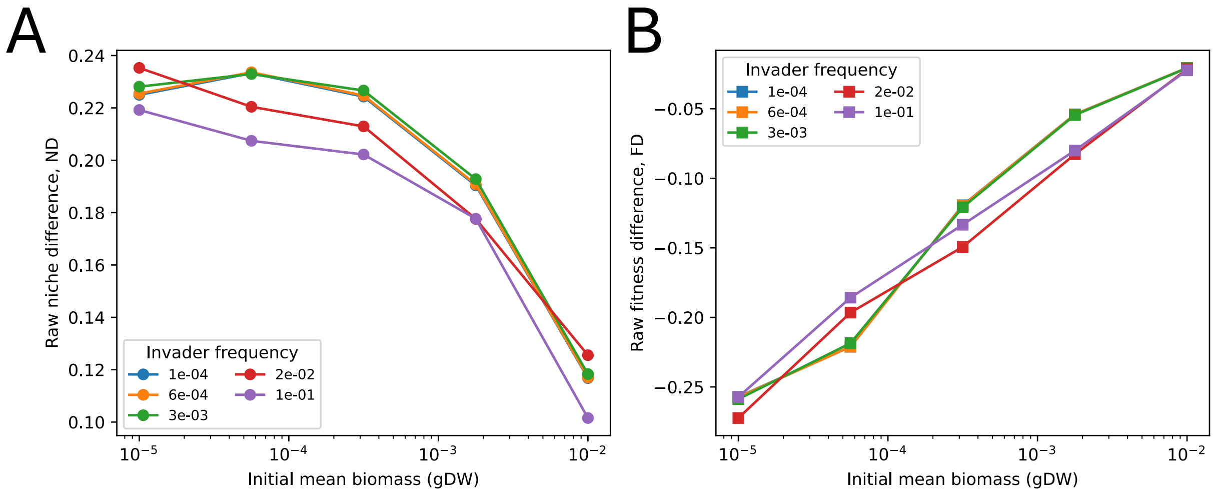


**Figure S7: Initial population size modulates estimated niche and fitness differences in batch dFBA while initial frequency does not.** **(A)** Raw niche difference and **(B)** raw fitness difference as functions of the initial *mean* biomass per strain (gDW) in reciprocal invasion assays. Colored curves correspond to different invader starting frequencies $p^{(L)} \in \{10^{-4}, 6 \times 10^{-4}, 3 \times 10^{-3}, 2 \times 10^{-2}, 10^{-1}\}$.

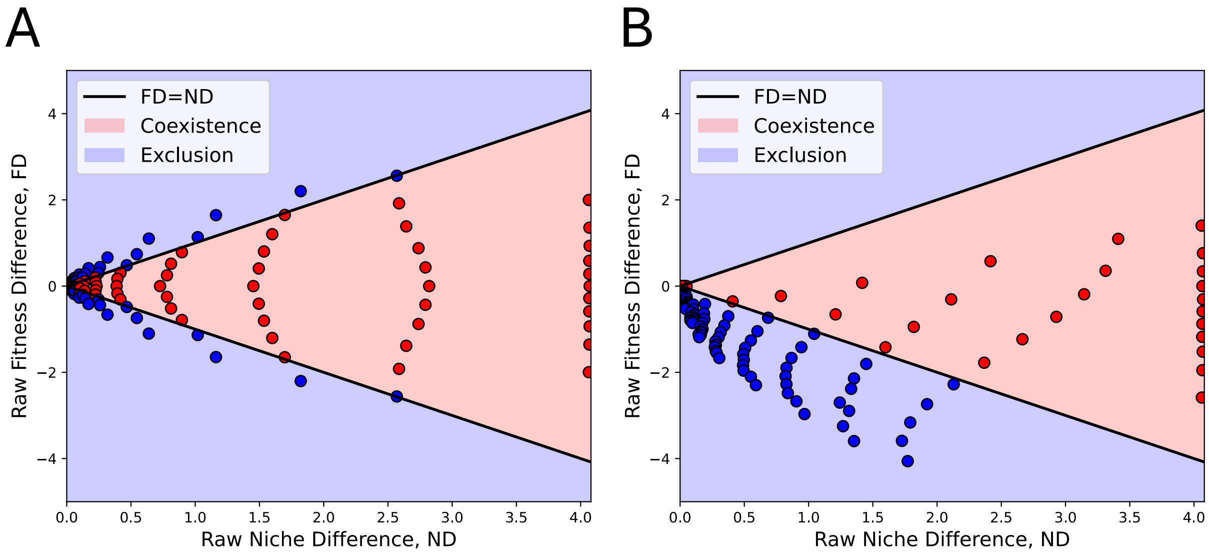


**Figure S8: Raw niche and fitness difference trajectories from neutrality.** **(A)** Plot corresponding to Figure 3A in the main text on raw (untransformed) niche and fitness difference axes. Trajectories departing from neutrality by changing the uptake flux, $T$, on alternate resources ($\beta$ = glucose, $\gamma$ = fructose) or resource concentration ratio $\frac{[C_{\text{glucose}}]}{[C_{\text{fructose}}]}$ from $\frac{5 \text{ mM glucose}}{50 \text{ mM fructose}}$ to $\frac{50 \text{ mM glucose}}{5 \text{ mM fructose}}$. **(B)** Plot corresponding to Figure 3B in the main text on raw (untransformed) niche and fitness difference axes. Trajectories from neutrality where $\beta =$ glucose and $\gamma =$ succinate by changing uptake flux on alternate resources or resource concentration ratios change as in (A).

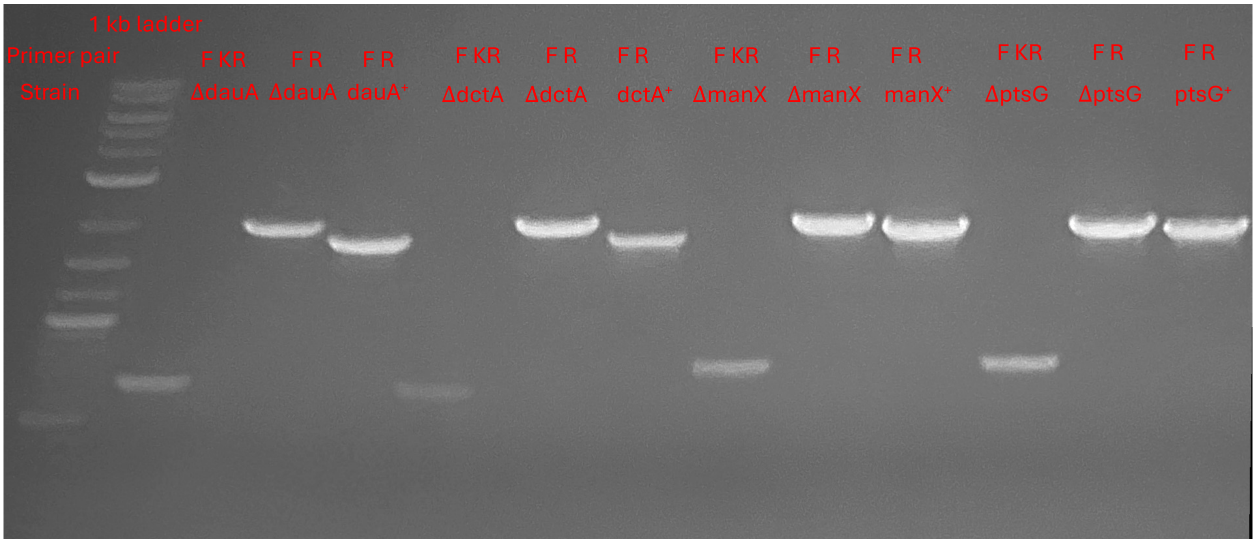


**Figure S9: Colony PCR confirmation of carbon transporter genotypes.** Agarose gel showing representative amplicons. Lanes are grouped in triplets by locus; within each triplet, the primer pair (upper label) and strain (lower label) are indicated. **F** is a gene-specific forward primer annealing at the 5′ end of the locus, in sequence retained in both alleles; **R** is a reverse primer just outside the 3′ end of the gene; **KR** is a reverse primer internal to the kan$^R$ cassette. The (F + KR) primers amplify the boundary between the chromosomal flank and the integrated cassette and therefore yield a product *only* when kan$^R$ has replaced the native ORF (deletion strains; first lane of each triplet). The (F + R) primers amplify across the locus: in the deletion strain the product spans the cassette-replaced allele, whereas in the corresponding $X^+$ strain it spans the intact ORF.

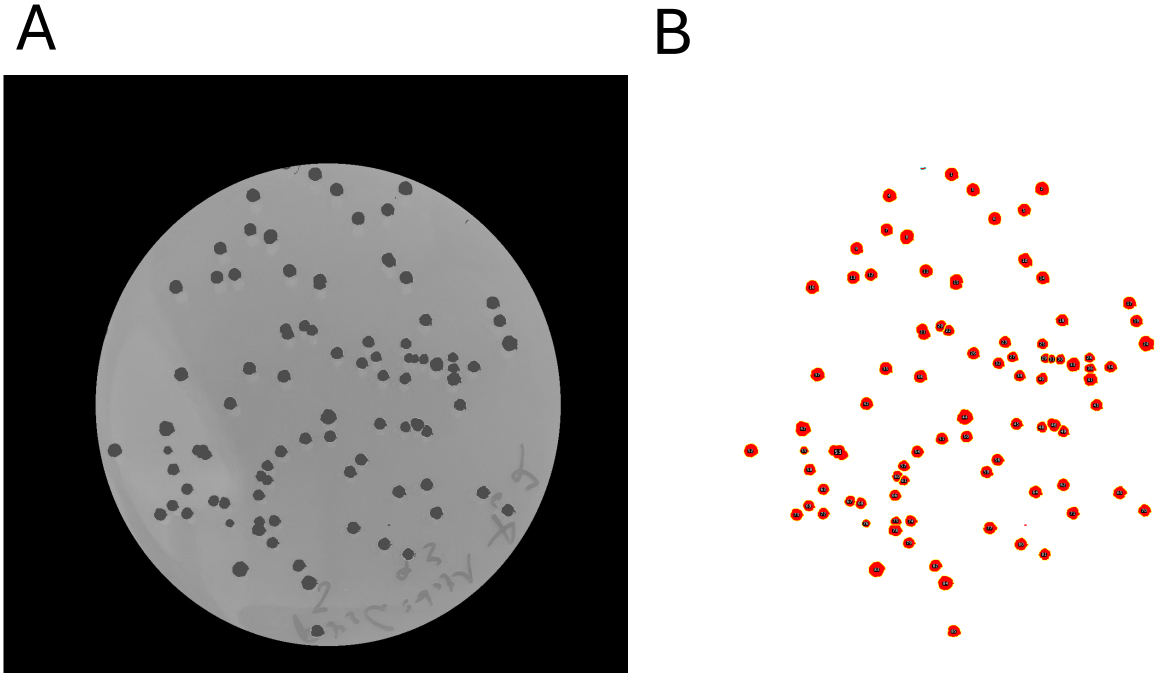

**Figure S10: Semi–automated colony counting in ImageJ. A** Cropped plate image after preprocessing: converted to 8-bit, rectangle crop to the plate, oval ROI to retain the usable agar. Colonies appear as dark spots against a uniform background (pre-threshold view shown). **B** Segmentation and enumeration overlay produced by the macro.

### Supplementary Tables

**Table S1:** Carbon sources and exchange reactions used in the dFBA simulations.

This table lists all carbon sources, their abbreviations, exchange IDs, chemical formulas, molecular weights, and the analysis set(s) in which they were used. Substrates labeled "Main text; Supp. A" were used in the focal coexistence simulations in the Main text and were also included in the extended single-substrate growth-rate predictor analysis in Supplementary Text A. Substrates labeled "Supp. A only" were used only in the extended growth-rate predictor analysis.

| Name | Abbrev. | Exchange ID | Formula | MW | Analysis set(s) |
|---|---|---|---|---|---|
| Glucose | Glc | EX_glc__D_e | $C_6H_{12}O_6$ | 180.16 | Main text; Supp. A |
| Fructose | Fru | EX_fru_e | $C_6H_{12}O_6$ | 180.16 | Main text; Supp. A |
| Succinate | Suc | EX_succ_e | $C_4H_6O_4$ | 118.09 | Main text; Supp. A |
| Galactose | Gal | EX_gal_e | $C_6H_{12}O_6$ | 180.16 | Main text; Supp. A |
| Glycerol | Glyc | EX_glyc_e | $C_3H_8O_3$ | 92.09 | Main text; Supp. A |
| D-Xylose | Xyl | EX_xyl__D_e | $C_5H_{10}O_5$ | 150.13 | Main text; Supp. A |
| Fumarate | Fum | EX_fum_e | $C_4H_4O_4$ | 116.07 | Main text; Supp. A |
| L-Malate | Mal | EX_mal__L_e | $C_4H_6O_5$ | 134.09 | Main text; Supp. A |
| Citrate | Cit | EX_cit_e | $C_6H_8O_7$ | 192.12 | Main text; Supp. A |
| Mannose | Man | EX_man_e | $C_6H_{12}O_6$ | 180.16 | Supp. A only |
| D-Ribose | Rib | EX_rib__D_e | $C_5H_{10}O_5$ | 150.13 | Supp. A only |
| D-Lactate | Lac | EX_lac__D_e | $C_3H_6O_3$ | 90.08 | Supp. A only |
| Trehalose | Tre | EX_tre_e | $C_{12}H_{22}O_{11}$ | 342.30 | Supp. A only |
| Maltose | Malt | EX_malt_e | $C_{12}H_{22}O_{11}$ | 342.30 | Supp. A only |
| D-Sorbitol | Sbt | EX_sbt__D_e | $C_6H_{14}O_6$ | 182.17 | Supp. A only |
| Acetate | Ac | EX_ac_e | $C_2H_4O_2$ | 60.05 | Supp. A only |
| Ethanol | EtOH | EX_etoh_e | $C_2H_6O$ | 46.07 | Supp. A only |
| Pyruvate | Pyr | EX_pyr_e | $C_3H_4O_3$ | 88.06 | Supp. A only |
| 2-Oxoglutarate | AKG | EX_akg_e | $C_5H_6O_5$ | 146.10 | Supp. A only |
| L-Alanine | Ala | EX_ala__L_e | $C_3H_7NO_2$ | 89.09 | Supp. A only |
| L-Aspartate | Asp | EX_asp__L_e | $C_4H_7NO_4$ | 133.10 | Supp. A only |
| L-Glutamate | Glu | EX_glu__L_e | $C_5H_9NO_4$ | 147.13 | Supp. A only |
| L-Serine | Ser | EX_ser__L_e | $C_3H_7NO_3$ | 105.09 | Supp. A only |

**Table S2:** Strain details for *E. coli* used in experimental assays.

This table lists the transporter genes deleted in each strain, along with a brief description of the knockout and its expected impact on resource utilization.

| Strain | Gene deletion | Kan$^R$ | Background strain | Notes |
|---|---|---|---|---|
| WT | None | No | MG1655 | Wildtype control |
| $\Delta nusB$ | *nusB* | Yes | MG1655 | Wildtype kan resistance (neutral). From SI Reference [3]. |
| $\Delta manX$ | *manX* | Yes | MG1655 | Glucose transporter deletion |
| $\Delta ptsG$ | *ptsG* | Yes | MG1655 | High-affinity glucose transporter deletion |
| $\Delta dauA$ | *dauA* | Yes | MG1655 | Succinate transporter deletion |
| $\Delta dctA$ | *dctA* | Yes | MG1655 | C4-dicarboxylate (succinate) transporter deletion |

**Table S3:** Primers used for colony PCR validation of transporter-gene deletions.

| Gene | Primer Name | Sequence (5'–3') | Purpose |
|---|---|---|---|
| *manX* | manX_F | CAAACGAATGTGACAAGGAT | Upstream |
| *manX* | manX_R | TGAAGAGTGGTAATCTCCAT | Control (gene) |
| *ptsG* | ptsG_F | GCCTTCAATCCATCCGTTGA | Upstream |
| *ptsG* | ptsG_R | GAAGGTTCTATCGTCTACGGC | Control (gene) |
| *dauA* | dauA_F | TCGAACCCGGCTCAAAGACCC | Upstream |
| *dauA* | dauA_R | CCGGAGGTGACATATGAAACG | Control (gene) |
| *dctA* | dctA_F | AGAGTTACCTGGAAGAAAAG | Upstream |
| *dctA* | dctA_R | GAATTTTTGTGCCCTGCAT | Control (gene) |
| *kanR* | KanR_R | CAATCCATCTTGTTCAATCAT | Insert |

**Table S4:** Best–fit maximum uptake bounds $\hat{T}$ ($\mathrm{mmol\,gDW^{-1}\,h^{-1}}$) from monoculture fitting, with RMSE of the fit.

| Strain | Substrate | $T$ | RMSE |
|---|---|---|---|
| WT | Glucose | 6.20 | 0.016 |
| WT | Succinate | 10.00 | 0.037 |
| $\Delta manX$ | Glucose | 7.40 | 0.012 |
| $\Delta manX$ | Succinate | 10.00 | 0.105 |
| $\Delta ptsG$ | Glucose | 2.67 | 0.018 |
| $\Delta ptsG$ | Succinate | 10.00 | 0.012 |
| $\Delta dauA$ | Glucose | 5.70 | 0.012 |
| $\Delta dauA$ | Succinate | 5.87 | 0.014 |
| $\Delta dctA$ | Glucose | 6.80 | 0.011 |
| $\Delta dctA$ | Succinate | 2.77 | 0.028 |

**Table S5:** Single-predictor regressions for maximum growth rate across the extended carbon source set. Each descriptor was fit in a separate linear model of the form $\mu \sim x_j$. Pearson $r$ gives the direction and strength of the association, $R^2$ summarizes in-sample explanatory power, and $R^2_{\text{LOOCV}}$ and LOOCV RMSE summarize leave-one-out predictive performance.

| Predictor | Category | Pearson $r$ | $R^2$ | RMSE | $R^2_{\text{LOOCV}}$ | LOOCV RMSE |
|---|---|---|---|---|---|---|
| $\text{NADH}_{prod}$ | Cofactor | 0.782 | 0.612 | 0.113 | 0.466 | 0.132 |
| $C_{\text{eff}}$ | Substrate | 0.757 | 0.573 | 0.119 | 0.465 | 0.133 |
| #C | Substrate | 0.680 | 0.463 | 0.133 | 0.175 | 0.165 |
| #active reactions | Network | 0.592 | 0.350 | 0.146 | 0.261 | 0.156 |
| $V_{\text{tot}}$ | Network | -0.277 | 0.077 | 0.174 | -0.157 | 0.195 |
| $\text{NADPH}_{prod}$ | Cofactor | -0.191 | 0.037 | 0.178 | -0.081 | 0.189 |
| $\text{ATP}_{prod}$ | Cofactor | 0.175 | 0.031 | 0.179 | -81.647 | 1.648 |
| Flux entropy $H_v$ | Network | -0.054 | 0.003 | 0.181 | -0.214 | 0.200 |
| Shortest path $d_s$ | Network/pathway | -0.018 | 0.000 | 0.181 | -0.135 | 0.193 |
| Bottleneck flux | Network/pathway | -0.011 | 0.000 | 0.181 | -0.098 | 0.190 |

Note: $R^2_{\text{LOOCV}} < 0$ indicates that leave-one-out predictions performed worse than predicting the mean $\mu$ for all substrates. The very poor LOOCV performance of $\text{ATP}_{prod}$ reflects sensitivity to a high-leverage substrate and was not interpreted as a robust predictor.

**Table S6:** Added-value regressions beyond NADH production for predicting maximum growth rate. The baseline model is $\mu \sim \text{NADH}_{prod}$. Each added-value model has the form $\mu \sim \text{NADH}_{prod} + x_j$. Negative ΔLOOCV RMSE and negative ΔAICc indicate improvement relative to the NADH-only model.

| Model / added predictor | Category | $R^2$ | $R^2_{\text{LOOCV}}$ | LOOCV RMSE | ΔLOOCV RMSE | ΔAICc |
|---|---|---|---|---|---|---|
| $\text{NADH}_{prod}$ only | Cofactor | 0.612 | 0.466 | 0.132 | – | – |
| #active reactions | Network | 0.701 | 0.594 | 0.115 | -0.017 | -2.994 |
| $C_{\text{eff}}$ | Substrate | 0.667 | 0.550 | 0.122 | -0.011 | -0.554 |
| Bottleneck flux | Network/pathway | 0.612 | 0.464 | 0.133 | 0.000 | 2.955 |
| Flux entropy, $H_v$ | Network | 0.636 | 0.441 | 0.136 | 0.003 | 1.511 |
| $\text{NADPH}_{prod}$ | Cofactor | 0.612 | 0.422 | 0.138 | 0.005 | 2.959 |
| Total flux, $V_{\text{tot}}$ | Network | 0.620 | 0.417 | 0.138 | 0.006 | 2.522 |
| Shortest path, $d_s$ | Network/pathway | 0.616 | 0.402 | 0.140 | 0.008 | 2.748 |
| #C | Substrate | 0.655 | 0.278 | 0.154 | 0.022 | 0.291 |
| $\text{ATP}_{prod}$ | Cofactor | 0.633 | 0.245 | 0.157 | 0.025 | 1.665 |
